# Heterogeneous therapy-resistant cancer cells have distinct and exploitable drug sensitivity profiles

**DOI:** 10.1101/2025.04.25.650475

**Authors:** Gianna T. Busch, Ryan H. Boe, Jingxin Li, Robert F. Gruener, Miles J. Arnett, Pavithran T. Ravindran, Meenhard Herlyn, R. Stephanie Huang, Arjun Raj

## Abstract

Resistance to targeted therapies is a significant clinical problem, but eliminating resistant cancer cells has proven difficult. One potential reason for this difficulty is heterogeneity in the resistant population: even genetically homogeneous cancer cell populations can give rise to many resistant subtypes, each potentially with specific second-line drug vulnerabilities. Using high-throughput drug screening of genetically-identical resistant clones with varying transcriptomes and morphologies, we show that each clone had a distinct drug sensitivity profile. These results suggested that there are drugs that are effective against only subsets of resistant populations but in combination eliminate a large proportion of the resistant population. Using the individual clone sensitivity profiles, we prospectively identified combinations that were highly effective at eliminating most of the resistant population. Our results demonstrate the effectiveness of “subpopulation-directed synergy”, showing that considering population heterogeneity can reveal therapeutic opportunities otherwise masked by population averages, offering new strategies to combat therapy resistance.

## INTRODUCTION

Melanoma is often driven by BRAF^V600E^ mutations that lead to constitutive activation of the MAPK pathway^1–4^. Inhibition of this mutant protein via targeted therapy initially causes many melanoma cells to die, but in the clinic, resistant populations of cells eventually emerge that are refractory to further treatments, leading to tumor recurrence and patient death^5–9^. Targeted therapies and immunotherapies can lead to cells that are cross resistant to each other, suggesting that strategies to combat therapy resistance could also be applicable to combating resistance to immunotherapies^10–17^. Strategies to target resistant cells thus have enormous therapeutic potential but have proven difficult to find.

One potential explanation for why therapy resistance has remained a significant problem is the heterogeneity of the resistant population. The fundamental challenge is the emerging realization that the resistant population is not a monolith, but rather is likely composed of several subpopulations of resistant cells, each of which may possess distinct therapeutic vulnerabilities^18^. The existence of different resistant “types” highlights the challenge of finding a second-line therapy: such a therapeutic must effectively target all these potentially diverse resistant cells. However, the existence of multiple resistant types also affords a new therapeutic opportunity. If one could identify drug vulnerabilities for each resistant type individually, then one could in principle identify combinations of drugs that would synergize at the population level to collectively eliminate a larger proportion of the resistant tumor cell population. For instance, it could be that Drug 1 kills one half of the resistant population while Drug 2 kills the other half (Fig. 1A). These drugs would not individually be considered viable second-line treatments, but together they could be highly effective. However, screens to evaluate all possible combinations would be prohibitively large. The advantage of screening individual resistant subpopulations is that, by focusing on resistant types that could have specific vulnerabilities rather than trying to find a single agent that could affect all resistant types, one might identify several novel drugs that may otherwise have been missed in a global screen.

**Figure 1.**
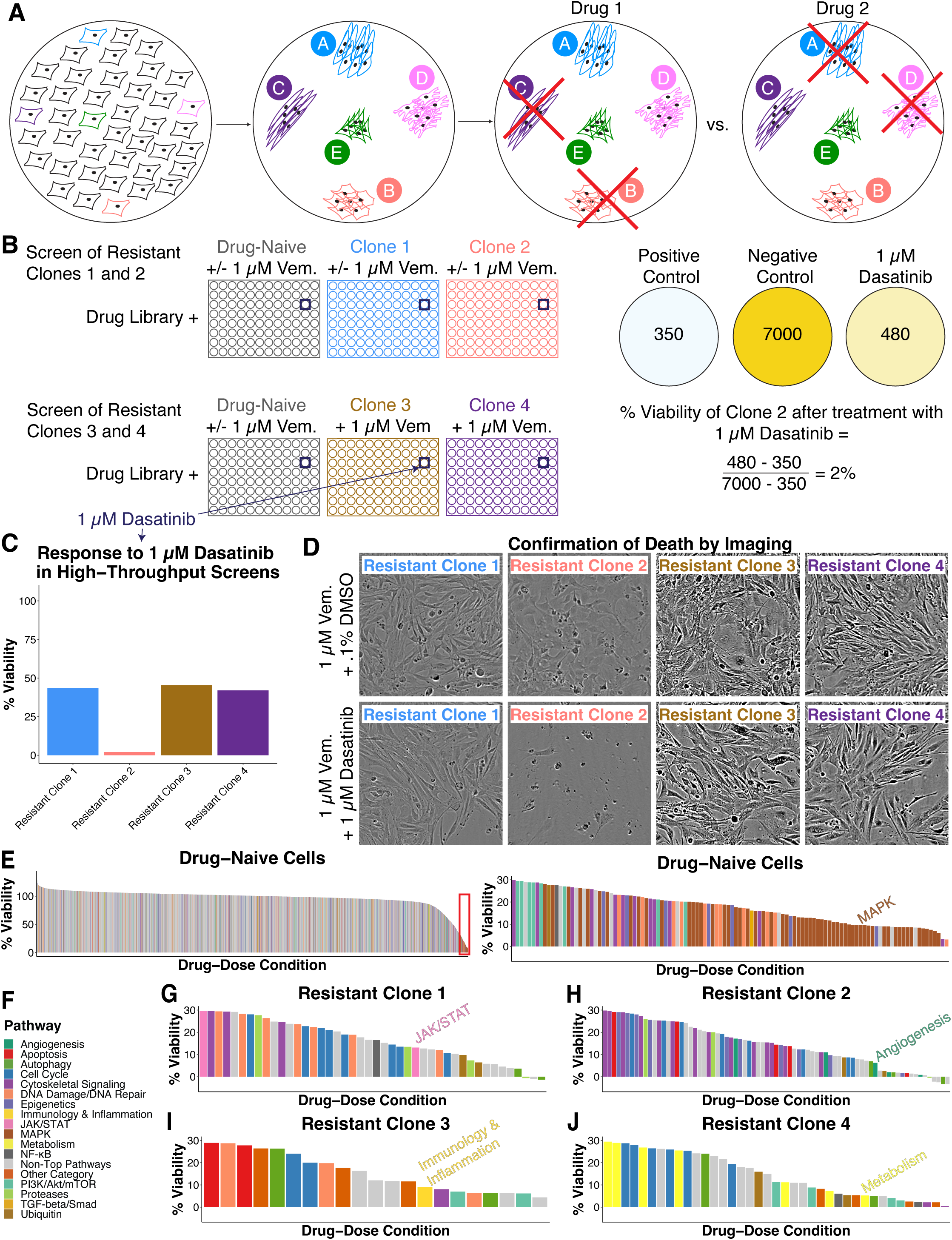
Genetically identical resistant clones have unique drug sensitivity profiles. A. Cartoon schematic showing the possibility of different resistant types being sensitive to different drugs. B. (Left) Schematic showing the composition of each high-throughput screen. Briefly, in the first screen, drug-naive melanoma cells and Resistant Clones 1 and 2 were screened using the pan-cancer drug library, both with and without 1 µM vemurafenib. In the second screen, drug-naive melanoma cells were screened using the library both with and without 1 µM vemurafenib, while Resistant Clones 3 and 4 were screened using the library only with 1 µM vemurafenib. (Right) Schematic showing how cell viability is calculated using the luminescence output of the CellTiter-Glo assay. C. Bar graph of the resistant clones’ responses to 1 µM dasatinib in the high throughput screens (seven days of drug treatment). The values are from screening with vemurafenib (n = 1). D. (Top row) Brightfield images of each resistant clone treated with 1 µM vemurafenib + 1 µM dasatinib for seven days. (Bottom row) Brightfield images of each resistant clone treated with 1 µM vemurafenib + .1% DMSO for seven days. E. (Left) Rank-order plot of the viability of all drug-dose conditions on drug-naive cells, excluding those at 10 µM and those which targeted (< 50% viability) all cell lines. (Right) Zoom-in of the remaining drug-dose conditions with < 30% viability. The top 7 most common pathways among the remaining drug-dose conditions with < 30% viability were calculated and used to color the plots. We chose a threshold of 50% as it passed the true difference threshold of 20 percentage points identified in our cross-batch analysis, so all remaining drug-dose conditions under 30% would satisfy this threshold. F. Pathway legend for all rank-order plots. G. Rank-order plot of the viability of drug-dose conditions with < 30% viability on Resistant Clone 1, excluding those at 10 µM and those which targeted (< 50% viability) all cell lines. The top 8 most common pathways among the remaining drug-dose conditions with < 30% viability were calculated and used to color the plot. We chose a threshold of 50% as it passed the true difference threshold of 20 percentage points identified in our cross-batch analysis, so all remaining drug-dose conditions under 30% would satisfy this threshold. H. Rank-order plot of the viability of drug-dose conditions with < 30% viability on Resistant Clone 2, excluding those at 10 µM and those which targeted (< 50% viability) all cell lines. The top 8 most common pathways among the remaining drug-dose conditions with < 30% viability were calculated and used to color the plot. We chose a threshold of 50% as it passed the true difference threshold of 20 percentage points identified in our cross-batch analysis, so all remaining drug-dose conditions under 30% would satisfy this threshold. I. Rank-order plot of the viability of drug-dose conditions with < 30% viability on Resistant Clone 3, excluding those at 10 µM and those which targeted (< 50% viability) all cell lines. The top 8 most common pathways among the remaining drug-dose conditions with < 30% viability were calculated and used to color the plot. We chose a threshold of 50% as it passed the true difference threshold of 20 percentage points identified in our cross-batch analysis, so all remaining drug-dose conditions under 30% would satisfy this threshold. J. Rank-order plot of the viability of drug-dose conditions with < 30% viability on Resistant Clone 4, excluding those at 10 µM and those which targeted (< 50% viability) all cell lines. The top 8 most common pathways among the remaining drug-dose conditions with < 30% viability were calculated and used to color the plot. We chose a threshold of 50% as it passed the true difference threshold of 20 percentage points identified in our cross-batch analysis, so all remaining drug-dose conditions under 30% would satisfy this threshold.

One potential explanation for the heterogeneity in the resistant population is the acquisition of distinct secondary resistance driver mutations, and it might thus be assumed that cataloguing all the genetic differences would be sufficient to identify all the secondary resistance mechanisms. However, recent work from our group has demonstrated that even a genetically homogeneous population of tumor cells can give rise to many different types of resistant cells, even under highly controlled settings. These genetically identical resistant cells showed marked differences in their transcriptome profiles, growth characteristics, and invasion phenotypes^19, 20^, suggesting that heterogeneity in resistant cell types is a fundamental problem not dependent on genetic variation, and we have demonstrated that these distinct resistant types appear in actual tumors^19, 21^. However, it remains unknown whether these distinct resistant types possess different drug sensitivity profiles that could be therapeutically exploited via subpopulation-directed synergy.

Here, we demonstrate that genetically identical resistant clones have distinct drug sensitivities. Using high-throughput drug screening of individual resistant clones derived from the same parental cell line, which maintain their distinct clonal identities over time and passaging, we show that each resistant clone exhibits its own distinct drug sensitivity profile to second-line inhibitors. Broad transcriptomic differences, beyond previously defined resistance programs, were predictive of sensitivity patterns among the clones. By identifying drugs that target different subsets of the resistant population, we prospectively designed combinations that effectively eliminated a larger proportion of resistant colonies than single agents alone. Our results demonstrate that considering population heterogeneity can reveal therapeutic opportunities otherwise masked by population averages, offering new strategies to combat therapy resistance.

## RESULTS

### Colony-derived cell lines enable high-throughput screening of different resistant types

Our recent work showed that individual therapy-resistant clones, even emerging from a genetically homogeneous initial population, can have stable differences in the transcriptome, morphology, proliferation rate, and invasiveness^19, 20^. These unique characteristics were highly consistent within a clone and maintained through multiple rounds of division and passaging^19^. We thus hypothesized that genetically identical resistant clones might also possess different sensitivities to second-line inhibitors (Fig. 1A).

To test this hypothesis, we established multiple clonal, resistant cell lines (resistant clones) derived from individual resistant colonies all emerging from the same parental cell line. We generated colonies by treating sparsely-plated clonal, drug-naive BRAF^VE^ melanoma cells with a targeted inhibitor or combination of targeted inhibitors. Once colonies reached sufficient size, we manually isolated and expanded each as an individual cell line, hereafter termed resistant clone (Supp. Fig. 1A, Methods). We isolated 43 resistant clones in total: 35 from treatment with 1 µM vemurafenib and eight from treatment with 250 nM dabrafenib/2.5 nM trametinib (Methods). Importantly, whole genome sequencing of multiple resistant clones (16 clones; all resistant to vemurafenib) revealed no recurrent mutations^19^. This finding corroborated that resistance was not driven by *de novo* genetic alterations, consistent with our prior results using fluctuation analysis (Methods)^19, 22^.

We used high-throughput drug screening to obtain a comprehensive picture of the drug sensitivity profiles among these resistant clones. To narrow the number of clones for screening, we reasoned that resistant clones with broad transcriptomic differences from each other would be most likely to show distinct drug sensitivity profiles. Of the 43 resistant clones we isolated, we performed bulk RNA sequencing on 30 and analyzed their expression of previously identified resistance programs (Methods)^19, 21^. Based on these transcriptome profiles, we selected four vemurafenib-resistant clones that represented diverse molecular phenotypes: Resistant Clone 1 (expressing the Smooth Muscle program), Resistant Clone 2 (expressing the canonical resistance marker *AXL)*^23^, Resistant Clone 3 (expressing the Extracellular Matrix program), and Resistant Clone 4 (expressing the Interferon program) (Supp. Fig. 1B).

We then screened these four resistant clones with a pan-cancer library containing 2240 drugs and small molecules with known pathways and targets against different cancer types. The screen was conducted by plating cells at a density of 1000 cells/well in 384-well plates and incubating with the library for seven days. As the library was spread out over multiple 384-well plates, we included a set of four drugs on every plate to serve as in-line validation; these were a drug that killed all cell lines (Epirubicin HCl), a drug that killed no cell lines (S3I-201), a drug to which Resistant Clone 1 was more sensitive (Clofarabine), and a drug to which Resistant Clone 2 was more sensitive (Dasatinib) (Supp. Fig. 2A). The readout was percent cell viability as measured by normalized CellTiter-Glo readings. CellTiter-Glo measures cell viability by generating a luminescent signal proportional to the amount of ATP present, which is directly proportional to the number of cells present in culture. We normalized the luminescent signal of each well using cells treated with the protease inhibitor Bortezomib to define the level of luminescence corresponding to the maximum amount of killing and cells treated with the vehicle DMSO to define the minimum amount of killing (Fig. 1B). Viability was defined as the fraction of total possible luminescence. We used a four-point dose curve of 0.01 µM, 0.1 µM, 1 µM, and 10 µM for each drug.

We performed the screen in two batches: Resistant Clones 1 and 2 in Screen A, and Resistant Clones 3 and 4 in Screen B, with the drug-naive cells in both (Fig. 1B). Note that in Screen A, we performed the screen for resistant and drug-naive cell lines both with and without vemurafenib, while in Screen B the resistant clones were screened together with vemurafenib and the drug-naive cells were screened both with and without vemurafenib (Fig. 1B). (We did not screen the resistant clones without vemurafenib in Screen B as analysis of Screen A data showed that the viability values of the library with and without vemurafenib were highly correlated, suggesting these conditions provided largely redundant information (Supp. Fig. 2B)).

To replicate the results of the screens in a different setting, we selected a set of eight drugs for follow-up testing. These drugs included the four control drugs noted above, as well as an additional four drugs of interest identified based on their differential sensitivity between Resistant Clones 1 and 2 (Supp. Fig. 2C). Seven of the eight drugs tested replicated the sensitivity profile of the high-throughput results (Fig. 1C-D, Supp. Fig. 2C). We observed via visual inspection that at a viability of 30%, as measured by CellTiter-Glo, the majority of cells on the dish appeared to be dead based on morphology (Supp. Fig. 2D); we thus considered 30% viability to correspond to an essentially complete elimination of the cells.

We first analyzed the viability of each drug across all resistant clones. We observed numerous drugs that appeared to differentially target either Resistant Clone 1 or 2 (Supp. Fig. 3A). We established appropriate statistical thresholds by using the drug sensitivities of drug-naive cells, which were included in both Screen A and B, as an across-batch test of differences arising from technical variability (Supp. Fig. 3B). For each drug, we measured the difference in viability between these replicates (Supp. Fig. 3C). Measuring the variance in this difference across all drugs, we adopted two standard deviations (representing an approximately 20 percentage point difference in viability) as our threshold for significance, which corresponded to an approximately 95% confidence interval (Supp. Fig. 3C-D). This confidence threshold was quite conservative because it also reflected batch differences between the two screens. We also used Screen A’s within-batch replicates (Resistant Clones 1 and 2 with and without vemurafenib) as another test of technical variability that did not reflect the batch differences (Supp. Fig. 3E-G). Doses of 0.01, 0.1, and 1 µM showed similar standard deviations across both the across-batch and within-batch comparisons. However, we observed significantly larger technical differences in cell viability between replicate screens at a dose of 10 µM; hence, we excluded this dose from subsequent analyses and experiments (Supp. Fig. 3C,F).

These high-throughput screens showed that individual resistant types have different drug sensitivity profiles.

### Resistant clones have differences in second-line inhibitor sensitivity

To determine whether individual resistant clones had different drug sensitivities, we sought to identify drugs that showed specific effects to particular resistant cell lines. We first found all drug-dose conditions to which at least one individual cell line was highly sensitive (viability < 30% in at least one line), then filtered out all drug-dose conditions to which all cells were sensitive (viability < 50% across all cell lines) (Supp. Fig. 4A). Our approach was conservative because it required an almost complete killing of at least one resistant cell line; there are likely other drug-dose conditions that have genuine differences between lines as well.

Of the 6720 possible drug-dose conditions (2240 drugs each screened at 3 different doses, excluding 10 µM), 167 reduced viability by more than 50% for all cell lines, including the drug-naive cells (Supp. Fig. 4B). Unsurprisingly, these drug-dose conditions largely targeted basic cellular functions such as division and metabolism (Supp. Fig. 4B). We filtered out these 167 generally killing conditions (Fig. 1E ( Left)) and ranked the viabilities of the remaining drug-dose conditions with viabilities below 30% for the drug-naive cells as shown in Fig. 1E (Right), as well as each of the resistant clones. By our threshold, we found 44 drug-dose conditions that showed specific effects for Resistant Clone 1, 77 for Clone 2, 21 for Clone 3, and 39 for Clone 4. These numbers indicate that there were multiple drugs whose effects were specific to particular resistant clones.

We wondered whether the drugs specific to each individual clone showed any enrichment for inhibition of any particular regulatory pathways. We calculated how often the pathway being inhibited by each drug was represented in each resistant clone’s subset of drug-dose conditions, and normalized this count to the total number of drugs in the panel inhibiting that pathway. We then ranked the top eight pathways for each cell line. As expected, the drug-naive cells showed MAPK pathway inhibitors as the top category, consistent with the known sensitivity of these melanoma cells to targeted therapies that inhibit various parts of the MAPK pathway (e.g. BRAF^V600E^ inhibitors like vemurafenib and dabrafenib, and MEK inhibitors like trametinib); drugs targeting this pathway made up 60 of the 112 drug-dose conditions below 30% viability for drug-naive cells (Fig. 1E (Right) - F). Also, resistant clones were not particularly sensitive to MAPK pathway inhibitors, as expected given they were selected to be resistant to these targeted therapies.

Each resistant clone, however, had a unique set of pathways represented among its top hits, both different from drug-naive cells and from each other. These were often supported by multiple drugs targeting said pathway, further supporting the functional role of the pathway. For instance, the top pathway unique to Resistant Clone 1 was JAK/STAT signaling, supported by the drugs AZ 960 and AT9283 (Fig. 1G). Both of the JAK/STAT-signaling inhibitors specifically inhibited the JAK2 protein.

Resistant Clone 2 showed unique enrichment in angiogenesis-, apoptosis-, and epigenetics-targeting drugs (Fig. 1H). Of particular interest was the Angiogenesis pathway due to the concordance of the specific targets of the drugs. Most were BCR-ABL, Src, or Syk family kinase inhibitors. Many of these inhibitors are known to have cross-reactivity with each other, suggesting they may all be targeting the same underlying mechanism in Resistant Clone 2^24–26^. In addition, most of the epigenetics-targeting drugs unique to Resistant Clone 2 were bromodomain and extraterminal domain (BET) inhibitors. This concordance strengthened the potential utility of these drugs against Resistant Clone 2.

Resistant Clones 3 and 4 had comparatively fewer drug-dose conditions in their subsets. However, they did show unique pathways among their top hits. Resistant Clone 3 had a hit targeting the transcription factor NRF2 in the Immunology & Inflammation pathway (Fig. 1I). Resistant Clone 4 had a number of hits targeting metabolic pathways (Fig. 1J).

Enrichment of various pathways (Ubiquitin, Proteases, Apoptosis, Epigenetics, NF-κB, Autophagy, Cell Cycle, DNA Damage/Repair, Cytoskeletal Signaling, and PI3K/Akt/mTOR) was shared across multiple or all resistant clones. However, we already filtered out all generally killing drug-dose conditions. Therefore, at least one other cell line was significantly less affected by the hits within these shared pathways. In addition, the specific targets represented were not always the same. For example, most of the PI3K/Akt/mTOR pathway inhibitors targeting Resistant Clone 2 were inhibiting the serine/threonine protein kinase GSK-3, whereas those targeting Resistant Clone 4 were inhibiting AMPK or Akt. In other cases, as with Autophagy inhibitors, the same drugs targeted the resistant clones, while the drug-naive cells were comparatively less sensitive. These findings further supported our hypothesis that genetically identical resistant clones can have distinct vulnerability profiles, even differing in which targets are most druggable within a particular signaling pathway, that could potentially be exploited for second-line treatments.

### Transcriptomic differences are associated with second-line inhibitor sensitivity patterns

Given that the different resistant clones did not have meaningful genetic differences between them but did have different transcriptomes, it was possible that these differences in gene expression were correlated with their drug sensitivity. For instance, we noticed that Resistant Clone 2 expressed the canonical resistance marker *AXL* at high levels, while the others did not^23^. Expression of *AXL* has been linked to susceptibility to dasatinib, to which Resistant Clone 2 was highly sensitive, at least raising the possibility that differences in gene expression could be correlated to differences in drug sensitivity in our resistant clones^27, 28^. We wondered whether such examples were just isolated associations or pointed to more systematic relationships.

Principal component analysis (PCA) of the transcriptome profiles of the cell lines in our study revealed that while principal component 1 separated drug-naive cells from all resistant clones, principal component 2 notably distinguished Resistant Clone 2 from the other resistant clones (Supp. Fig. 5A). Similarly, PCA of sensitivity profiles showed comparable clustering patterns, with Resistant Clone 2 again separating from the other clones (Supp. Fig. 5B).

To quantify the correlation between drug sensitivity and transcriptome, we calculated distances between each pair of resistant clones in both transcriptomic and drug sensitivity PCA space. We found a strong correlation between these two sets of distances (Pearson r = 0.817, Mantel p = 0.042), indicating that clones with more similar transcriptomes exhibited more similar drug responses (Supp. Fig. 5C)^29^. Importantly, this correlation persisted even when Resistant Clone 2 was excluded from the analysis (Supp. Fig. 5C), suggesting a robust relationship between transcriptomic profiles and drug sensitivity patterns.

We have previously shown that the transcriptomes of individual resistant cells could be organized into different recurring resistance programs (meaning groups of genes whose expression was associated with particular resistance types). These resistant programs appeared in both *in vitro* and in patient samples ^19, 21^. We wondered whether these resistance programs might be more or less closely associated with drug sensitivity profiles than the PCA-based analysis of the transcriptome above. We quantified differential sensitivity between resistant clones across multiple drug-dose conditions, comparing these differences to established confidence thresholds. Resistant Clone 2, which exhibited the most distinct transcriptomic profile, also showed significantly more differential drug responses when compared to Resistant Clones 3 and 4, but comparisons between Resistant Clones 1, 3, and 4 revealed minimal differences in drug sensitivity profiles despite these clones expressing different resistance programs (Supplementary Figure 6D-E). These findings suggest that broader transcriptomic differences, rather than specific resistance programs, more accurately predict patterns of drug sensitivity in resistant clones.

### Subsets of resistant clones share some drug sensitivities

Our high throughput screening results suggested that individual resistant clones were highly variable in their response to second-line inhibitors. It remained unclear whether the clones profiled represented distinct archetypes, or if every clone had a unique drug sensitivity profile. However, conducting screens on a large number of cell lines was prohibitive. Therefore, we decided to curate a smaller panel of 24 drugs for testing across this broader swath of cell lines.

We wanted to focus our search on drugs that represented maximally different effects among the clones where one or more were dead and the remainder were virtually unaffected. We thus attempted to categorize all 6720 drug-dose conditions as universal, shared, or unique killers.

We defined universal killers as killing all clones, shared killers as killing multiple clones and not killing at least one other clone, and unique killers as killing only one clone and not killing at least one other clone. To maximize the degree of specificity, we used a threshold of < 30% viability for a drug-dose condition to kill a clone and > 70% to leave a clone alive.

For example, the KLF5 inhibitor SR18662 was a unique killer at a dose of 1 µM as after treatment with it Resistant Clone 4 had a viability of less than 30%, Resistant Clone 1 had a viability of approximately 50%, and Resistant Clones 2 and 3 had viabilities above 70% (Supp. Fig. 6A). Meanwhile, the PI3K/Akt inhibitor Deguelin was a shared killer at a dose of 1 µM as after treatment with it Resistant Clones 3 and 4 had viabilities of less than 30%, but Resistant Clones 1 and 2 had viabilties greater than 70% (Supp. Fig. 6B).

Although some drug-dose conditions were universal across all cell lines (66 total universal killers; often targeting general cellular pathways), more were specific to individual resistant clones (107 total unique killers) or shared among several clones (21 total shared killers). Unique and Shared killers spanned a variety of pathways, and all cell lines had at least one drug that uniquely killed it (Fig. 2A-B)^30^. Universal killers were largely related to basic cell functions such as cell cycle, metabolism, and protease function (Fig. 2C). We selected a set of 24 drugs for our panel—including the four control drugs we used in the screens—that covered unique killers for each clone, shared killers for different pairs of clones, and drugs that were shared killers for all of the resistant clones but not the drug-naive cells (thus eliminating general cell-killing agents) (Supp. Table 2).

**Figure 2.**
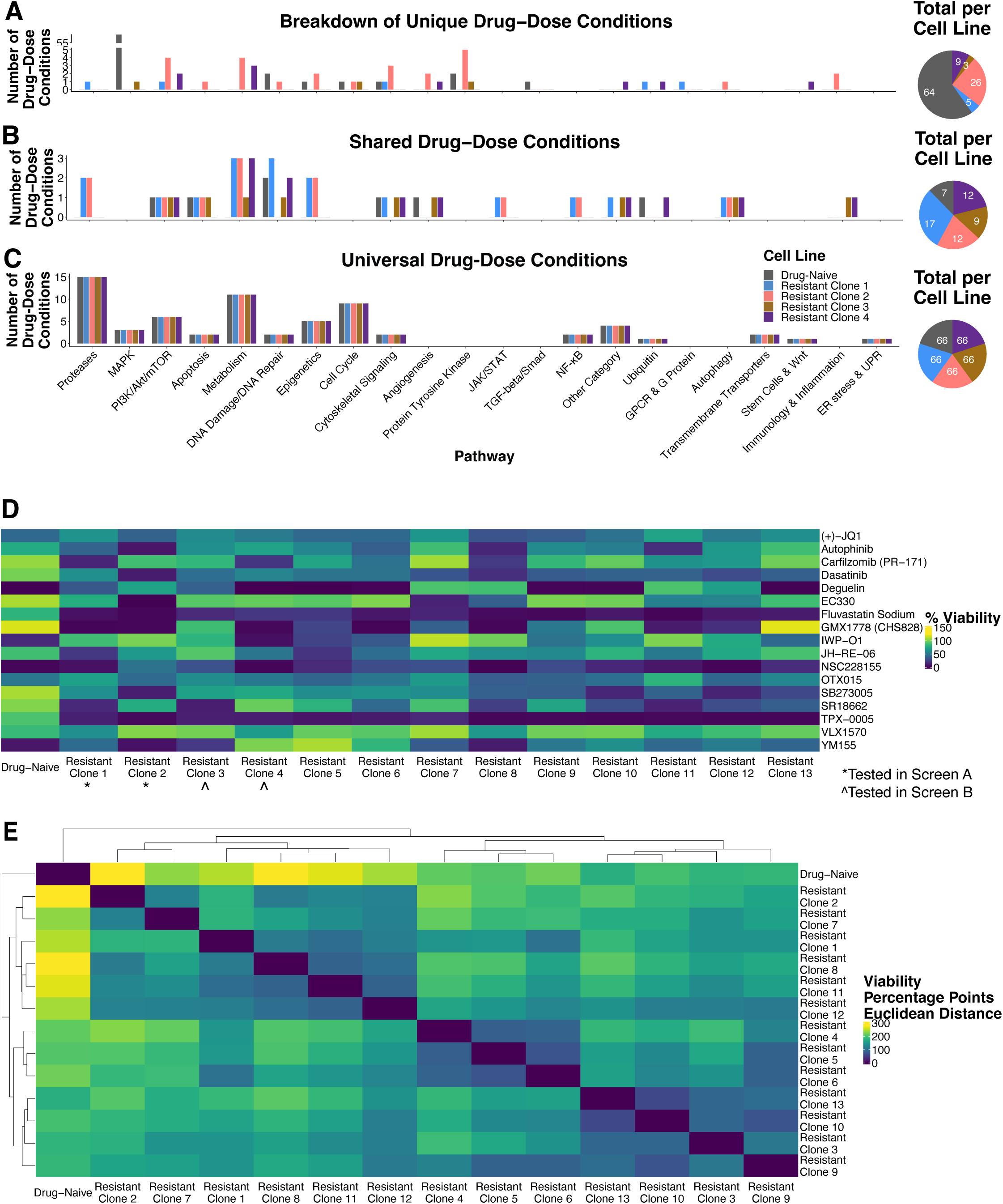
Certain drug sensitivity profiles may be shared across resistant clones. A. Bar plot of the number of drug-dose conditions that uniquely target a particular cell line, grouped by pathway. Each resistant cell line has a unique sensitivity fingerprint. Uniquely targeting is defined quantitatively as a drug-dose condition that has less than 30% viability for that particular cell line and greater than 70% viability for at least one other cell line. The pie chart shows the total for each cell line. B. Bar plot of the number of drug-dose conditions that target multiple cell lines, but not all, grouped by pathway. Shared targeting is defined quantitatively as a drug-dose condition that has less than 30% viability for multiple cell lines and greater than 70% viability for at least one other cell line. The pie chart shows the total for each cell line. C. Bar plot of the number of drug-dose conditions that target all cell lines, grouped by pathway. Universal targeting is defined quantitatively as a drug-dose condition that has less than 30% viability for all cell lines. The pie chart shows the total for each cell line. D. Heatmap of the viability data from Follow-Up Panel A. The drugs that did not validate based on the high-throughput screening results are excluded. Each drug is shown at the dose of interest as identified by the high-throughput screening data. All conditions were tested in duplicate, and the average of those two values is shown (n = 2). E. Heatmap of the Euclidean distance matrix of the data shown in E, clustered by rows and columns.

Next, we selected a set of 13 resistant clones to test with our follow-up panel along with the drug-naive cells (Supp. Table 3). This set included the four vemurafenib-resistant clones used in our high-throughput screening (Resistant Clones 1 - 4), five additional clones resistant to vemurafenib (Resistant Clones 5 - 9), and four clones generated via combination dabrafenib/trametinib treatment (Resistant Clones 10 - 13). Resistant clones were screened in media containing either vemurafenib or dabrafenib/trametinib, depending on their primary resistance. Drug-naive cells were screened without any additional inhibitor (only the vehicle DMSO).

The follow-up panel was conducted in one phase. Cells were plated at a density of 1000 cells/well in 384-well plates and incubated with the panel for seven days. The CellTiter-Glo Assay was then used to measure cell viability, and the readout was normalized to a positive control value (the protease inhibitor Bortezomib) and a negative control value (the vehicle DMSO) (Fig. 1B). Two technical replicates were used for this panel, and we observed high concordance between them (Supp. Fig. 7A).

An 8-point logarithmic dose curve was used, with the highest dose varying based on the dose at which we had seen differences among the resistant lines in the high-throughput screens (Supp. Table 2). We varied the starting dose in this way because we hypothesized that drugs showing stark differences at lower doses and not higher doses might also have interesting differences at even lower doses. Twenty of the drugs (including all four of the control drugs) were validated based on the results seen in the high-throughput screens and validation experiments and were included in the analysis (Supp. Fig. 7B).

If the resistant clones profiled in the high-throughput screens represented sensitivity archetypes, then we would expect the additional clones tested in our follow-up panel to resemble one of our original clones in terms of their sensitivity profiles. We generated a heatmap of the viability values for each cell line, showing only the dose of each drug where we observed the cell lines to be maximally different (Fig. 2D, Supp. Fig. 7C). Although we saw that each clone had some differences from the others, we wanted to quantify these differences more quantitatively, taking into account the entirety of the profile. We calculated the Euclidean distance between each pair of cell lines (all resistant clones and the drug-naive cells) (Fig. 2E). Our analysis showed some evidence for sensitivity archetypes, although there are still many unique sensitivities even within these archetypes.

We wondered whether the primary treatment to which the clones were resistant (vemurafenib or dabrafenib/trametinib) determined their sensitivity profiles. We found that resistant clones derived from colonies resistant to primary dabrafenib/trametinib treatment (Resistant Clones 10 - 13) did not cluster separately from the vemurafenib-resistant clones, indicating that sensitivity profiles are independent of the specific primary inhibitor(s) used in initial treatment.. Previous work has shown that the same transcriptomic types emerge whether resistant cells were treated with vemurafenib or combination dabrafenib/trametinib^19, 21^. Together, these results demonstrate that neither the transcriptomic type nor the sensitivity profile depend on the primary inhibitor, suggesting that the resistant types may be somewhat universal.

To quantify the extent of similarity and difference between resistant clones, we examined the most similar and most different clone pairs based on their Euclidean distances. Resistant Clones 10 and 13 showed remarkable similarity, exhibiting the smallest Euclidean distance among all resistant pairs. Their sensitivity profiles were so closely aligned that none of the drug-dose conditions differed by more than 40 percentage points in our distance analysis, with only three conditions exceeding our 20-point confidence threshold. In contrast, Resistant Clones 2 and 4, which represented the maximum observed Euclidean distance, differed by more than 40 percentage points across nine drug-dose conditions, with one additional condition surpassing our confidence threshold. Overall, the clusters we observed suggest that there may be certain combinations of sensitivities and invulnerabilities to different drug-dose conditions that are shared across multiple resistant clones, rather than entirely novel profiles for each clone.

### Drug sensitivities of resistant clones predict colony elimination patterns in larger resistant populations

Our high-throughput and targeted screening results identified distinct drug sensitivity profiles among genetically identical resistant clones. However, the resistant clones we used in high-throughput screening were generated by isolating and expanding only a small subset of colonies, which we have shown are biased towards those that grow suitably for large-scale experiments, and thus may represent only a subset of the resistant population^19^. Therefore, we next asked whether the sensitivity patterns we observed would be recapitulated with the full complement of resistant colonies. We specifically wondered whether drugs with clone-specific effects—those that strongly affected some resistant clones while leaving others completely unaffected—would show a bimodal elimination pattern with complete eradication of susceptible subpopulations and no effect on others, while drugs with moderate to strong effects across all clones would eliminate the majority of the resistant population. Killing some resistant colonies while leaving others undisturbed would support our initial premise that heterogeneous resistant populations could be targeted through “subpopulation-directed synergy,” where drugs that individually target distinct subsets of resistant cells.

CellTiter-Glo measures cell viability across the entire population, and so it is not well-suited to measure the effects of drugs on particular subpopulations, which requires single colony-resolution. We set up a colony formation assay to test the effects across the full spectrum of emergent resistant types (Supp. Fig. 8A). We plated drug-naive cells and treated them with 1 µM vemurafenib, resulting in the survival and colony formation of about 1 in 1000 cells. After three weeks, we then either continued treatment with vemurafenib only or added a second drug for two additional weeks. We monitored and quantified colony formation for all conditions at three, four, and five weeks after the initial treatment.

We selected drugs for testing based on the two distinct sensitivity patterns observed in our high-throughput screening data, drugs with specific effects and drugs with broad moderate to strong effects, and tested each at the lowest dose at which we observed the differential sensitivity pattern. For drugs with specific effects, we chose SR18662, a KLF5 inhibitor that preferentially targeted Resistant Clone 3, with a moderate effect on Resistant Clone 1, a small effect on Resistant Clone 2, and little to no effect on Resistant Clone 4 or the drug-naive cells at lower doses (Supp. Fig. 6A). We also included the Src inhibitor saracatinib that preferentially targeted Resistant Clone 2 with minimal effect on other resistant clones or drug-naive cells (Supp. Fig. 8B). For the moderate to strong effect category, we selected dasatinib, a BCR-ABL inhibitor that strongly targeted Resistant Clone 2 while exerting moderate effects on the other three resistant lines (Supp. Fig. 8C). Similarly, the bromodomain inhibitor (+)-JQ1 showed a moderate reduction in cell viability across all resistant lines (Supp. Fig. 8D). As a negative control, we included the STAT inhibitor S3I-201, which had no effect on any resistant clones or drug-naive cells (Supp. Fig. 8E).

Addition of SR18662 (1 μM) after 21 days of vemurafenib treatment resulted in approximately a 60% reduction in colony numbers compared to vemurafenib-only treatment at the final 5-week timepoint (Supp. Fig. 8G). Saracatinib (1 μM) resulted in approximately a 50% reduction in colony numbers at 5 weeks when added after 21 days of vemurafenib treatment (Supp. Fig. 8G). Although the specific effect of saracatinib was recapitulated, this effect was surprising given that we used a lower dose (1 μM) than where we observed differential effects among resistant clones (10 μM) to minimize off-target effects (Supp. Fig. 8B,H). None of the resistant clones were killed at a dose of 1 μM saracatinib (Supp. Fig. 8G). This replication of saracatinib’s specific effect at a lower dose suggests that some drugs with specific effects may be more effective at lower doses against the resistant colonies as compared to the resistant clones, potentially due to increased stability of clones from longer passaging durations. In contrast, the addition of Dasatinib (0.1 μM) killed most of the colonies by 5 weeks (Supp. Fig. 8I), and (+)-JQ1 (0.1 μM) also showed colony-killing activity at 5 weeks, though somewhat less than dasatinib (Supp. Fig. 8J). As expected, the negative control S3I-201 (1 μM) did not kill any of the colonies (Supp. Fig. 8K).

Our results showed that a drug’s effect on resistant clones in our high-throughput screens generally predicted its ability to kill the full slate of resistant colonies. SR18662’s specific effect on the resistant clones was recapitulated when it was added to resistant colonies. In contrast, dasatinib and (+)-JQ1 had moderate to strong effects on all resistant clones, and both killed most, although not all, of the resistant colonies. These findings validated our approach to using both high-throughput and targeted screening of select resistant clones to expand the search space of potential second-line therapies to treat the full spectrum of therapy-resistant cancer cells.

### Combinations of drugs can eliminate the resistant population

Given that the resistant clones’ unique drug sensitivities were recapitulated across the full spectrum of colonies, we hypothesized that combinations of drugs targeting different subsets of resistant clones could effectively eliminate the entire resistant population (Fig. 3A). Our approach of using both high-throughput and targeted screening of select resistant clones potentially identified new drugs that kill subsets of resistant cells, so we wanted to test whether they actually work in combination to eliminate more of the resistant population.

**Figure 3.**
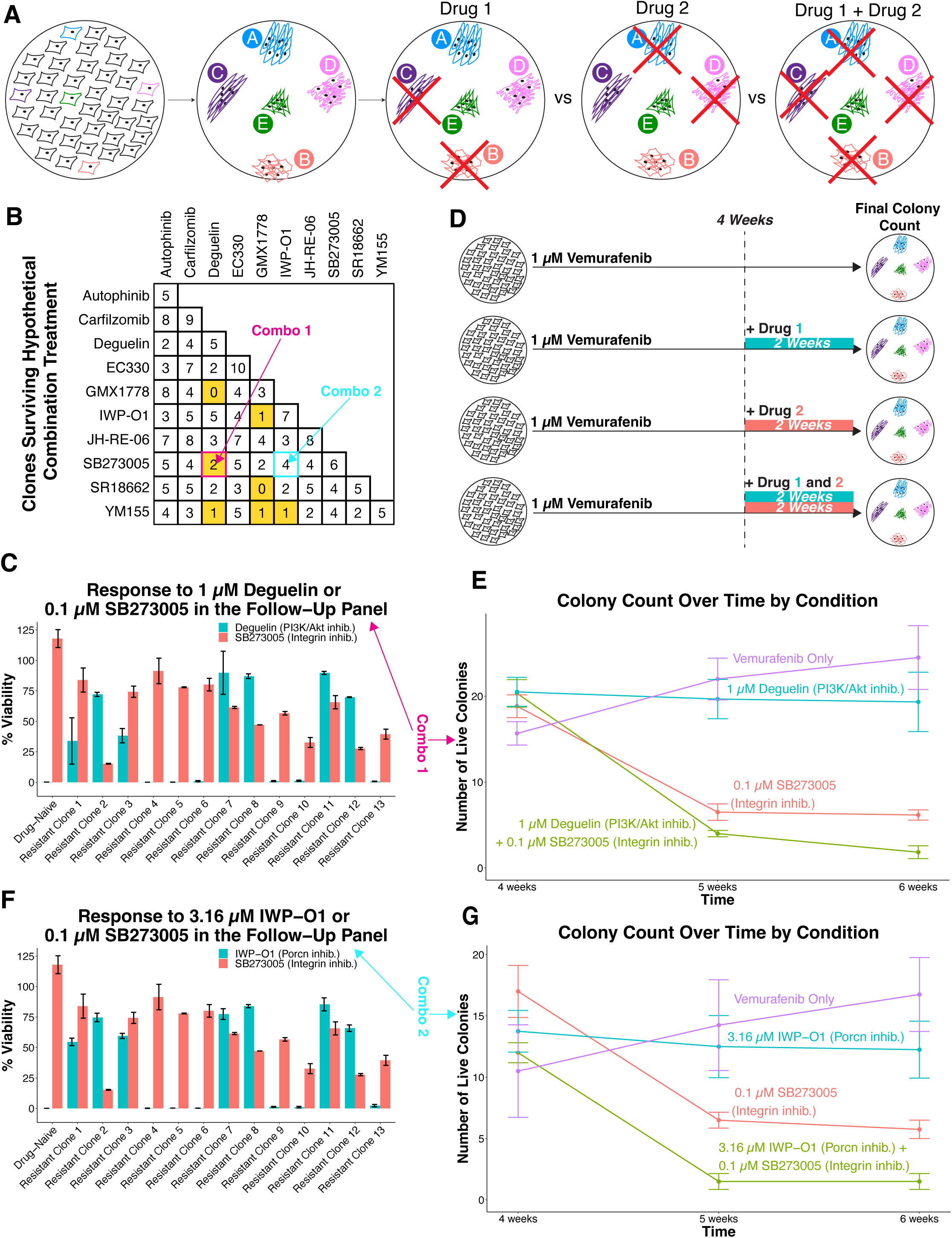
Combinations of secondary drugs can eliminate resistance. A. Cartoon schematic showing the possibility of different resistant types being sensitive to different drugs, with the combination of these drugs being able to eliminate a greater fraction of the resistant population than either as a single agent. B. Chart showing the number of resistant clones left alive after hypothetical combination treatment with pairs of drugs used in Follow-Up Panel A. Combinations which were tested are filled in yellow. C. Paired bar graph of the drug-naive cells and resistant clones’ responses to 1 µM deguelin or 0.1 µM SB273005 in the follow-up panel (seven days of drug treatment). The values for the resistant clones are from screening with vemurafenib (Clones 1 - 9) or dabrafenib/trametinib (Clones 10 - 13). The value for the drug-naive cells is from screening without vemurafenib. All conditions were tested in duplicate, and the average of the value is shown (n = 2). D. Cartoon schematic showing the experimental design for the combination experiments. Briefly, drug-naive cells were plated and treated with vemurafenib for four weeks to allow colonies to form. After four weeks, the colonies were subjected to one of four conditions: vemurafenib only for two weeks, the addition of one drug for two weeks, the addition of another drug for two weeks, or the addition of both drugs for two weeks. Colony counts were quantified and compared at four, five, and six weeks for all conditions. E. Line graph of the number of live colonies at four, five, and six weeks after treatment with vemurafenib only, vemurafenib with deguelin added after four weeks, vemurafenib with SB273005 added after four weeks, or vemurafenib with both drugs added after four weeks. The number of live colonies is the average across three experiments, each of which was done in duplicate (n = 6). F. Paired bar graph of the drug-naive cells and resistant clones’ responses to 3.16 µM IWP-O1 or 0.1 µM SB273005 in the follow-up panel (seven days of drug treatment). The values for the resistant clones are from screening with vemurafenib (Clones 1 - 9) or dabrafenib/trametinib (Clones 10 - 13). The value for the drug-naive cells is from screening without vemurafenib. All conditions were tested in duplicate, and the average of the value is shown (n = 2). G. Line graph of the number of live colonies at four, five, and six weeks after treatment with vemurafenib only, vemurafenib with IWP-O1 added after four weeks, vemurafenib with SB273005 added after four weeks, or vemurafenib with both drugs added after four weeks. The number of live colonies is the average across two experiments, each of which was done in duplicate (n = 4).

We used the viability data from our follow-up panel (Fig. 2E) to identify combinations of drugs that would kill distinct subsets of the resistant clones and therefore together theoretically kill all or nearly all clones. (Here, we used a more lenient threshold for killing to help us identify more potential synergies; see Methods.) We totaled the number of resistant clones killed by each drug and calculated the theoretical total of resistant clones killed by each combination of drugs, assuming independent effects with no double-counting of clones affected by both drugs (Fig. 3B, Supp. Fig. 9). This analysis identified several promising candidate drug combinations, for instance, the PI3K/Akt inhibitor Deguelin and the integrin inhibitor SB273005. These drugs showed particularly complementary coverage: Deguelin effectively killed one subset of resistant clones while SB273005 killed a different subset, with only two clones showing greater than 50% viability after treatment with either drug (Fig. 3C).

To test which combinations of drugs were effective in vitro, we used a colony formation assay with multiple treatment conditions in which secondary drugs were added at different times relative to vemurafenib (Fig. 3D). We first plated drug-naive cells and initiated treatment with 1 µM vemurafenib. We then created several experimental arms: (1) vemurafenib alone for the full six weeks as a control; (2-3) vemurafenib followed by the addition of either of two secondary drugs after four weeks; and (4) vemurafenib followed by the addition of both second drugs after four weeks. We monitored and quantified colony formation for all conditions at four, five, and six weeks after the initial treatment.

The addition of Deguelin (1 μM) after four weeks of vemurafenib treatment resulted in approximately a 20% reduction in the final number of colonies compared to treatment with vemurafenib only, which was lower than we would have expected based on our panel data (Fig. 3E). The addition of SB273005 (0.1 μM) after four weeks of vemurafenib treatment resulted in approximately a 66% reduction in the final number of colonies compared to treatment with vemurafenib only, which was slightly higher than we expected (Fig. 3E). Nevertheless, the addition of both of these drugs resulted in the death of all or one to two resistant colonies (Fig. 3E), confirming our hypothesis that combinations targeting complementary subsets of resistant cells could achieve more comprehensive elimination than either drug alone.

Next, we tested the combination of SB273005 (0.1 μM) and the porcn inhibitor IWP-O1, which hypothetically would kill all but four of the resistant clones (Fig. 3B, F). The addition of IWP-O1 (3.16 μM) after four weeks of vemurafenib treatment resulted in approximately a 33% reduction in the final number of colonies compared to treatment with vemurafenib only (Fig. 3G). As shown in panel E, addition of SB273005 (0.1 μM) after four weeks of vemurafenib treatment resulted in approximately a 66% reduction in the final number of colonies compared to treatment with vemurafenib only (Fig. 3G). Unexpectedly, the addition of both of these drugs resulted in the death of all or one to two resistant colonies, likely due to the stronger than average expected of SB273005 (Fig. 3G). The combinatorial effect of SB273005 and IWP-O1 further supports our hypothesis that combinations of drugs could target distinct fractions of resistant cells to achieve more comprehensive elimination.

We also tested the combination of the survivin inhibitor YM155 and the NAMPT inhibitor GMX1778, which hypothetically would kill all but one of the resistant clones (Fig. 3B). Since each drug individually was effective against a large fraction of clones, we reduced the dosage to target distinct subsets with greater specificity (Supp. Fig. 10A-B). The addition of YM155 (5 nM) after four weeks of vemurafenib treatment resulted in a 66% reduction in the final number of colonies compared to vemurafenib alone (Supp. Fig. 10C). In contrast, we found that the addition of GMX1778 (5 nM) after four weeks of vemurafenib treatment resulted in the death of all resistant colonies by six weeks, whether this drug was added alone or in combination with another drug (Supp. Fig. 10C). Nevertheless, a fraction of the colonies were still alive at five weeks, and we observed that the combination of GMX1778 and YM155 was more effective at this timepoint than either drug alone, eliminating approximately twice as many colonies as GMX1778 by itself (Supp. Fig. 10C). We also observed this behavior when GMX1778 was combined with each of three additional drugs, with the combination being more effective at five weeks than GMX1778 alone, although treatment with GMX1778 killed all resistant colonies by six weeks (Supp. Fig. 10D-F).

Finally we tested the combination of Deguelin and IWP-O1. This combination hypothetically would not kill all of the resistant clones (Fig. 3B). Indeed, the combination of Deguelin (1 μM) and IWP-O1 (3.16 μM) was not more effective than either drug as a single agent (Supp. Fig. 11A), demonstrating that drugs must be targeting complementary sets of resistant cells to achieve more complete elimination in combination.

We tested the remaining combinations identified as promising based on our follow-up panel viability data and found that the combinations did not outperform either drug as a single agent (Supp. Fig. 11B-D). Together, our results served to prove the principle that drugs that may not have been viable individually could indeed target different subsets of colonies and so, in combination, could eliminate a larger percentage of the resistant population.

### IDACombo accurately predicts combination efficacy

Our initial approach to identifying drug combinations targeting distinct resistant subpopulations yielded mixed results, with only one of our initial eight hypothesized drug pairs demonstrating improved combination efficacy. We thus used IDACombo, a computational method that predicts combination efficacy based on single-agent therapy response data^31, 32^. IDACombo operates under the assumption of independent drug action (IDA), which posits that the effectiveness of a drug combination on a particular cell line is determined by the most effective single drug in that combination, without synergistic effects^31–35^. Under that assumption, it then predicts how effective a combination would be across a large number of cell lines. It has been shown that IDACombo alone (without additivity or synergy) is enough to explain the benefit for many FDA approved drug combinations^34^. Given that the goal of IDACombo was to tackle population-level heterogeneity, the principle underlying it aligns well with our subpopulation-directed approach of targeting intrinsically heterogeneous resistant populations with drugs selected based on the sensitivity profiles of individual resistant clones.

We applied this method to all pairwise combinations in our follow-up panel to predict their effectiveness across our collection of resistant clones (Methods). Briefly, we calculated the IDACombo score for each pair of drugs at each available dose and averaged the score over the available dose range to generate an averaged score for each drug pair (Supp. Fig. 12A). Averaging IDACombo scores over all doses ensures the IDACombo effect is robust and seen at multiple doses. However, since the number of doses a drug has has an impact on viability changes for every drug, this would also influence the average IDACombo scores. We refined our analysis to ensure the best candidate drug combinations were still the best when correcting for this by averaging only IDACombo scores at “effective drug concentrations”, with “effective drug concentrations” meaning any monotherapy concentration for a drug that lowered the average viability of the cells to at least 90% (Fig. 4).

**Figure 4.**
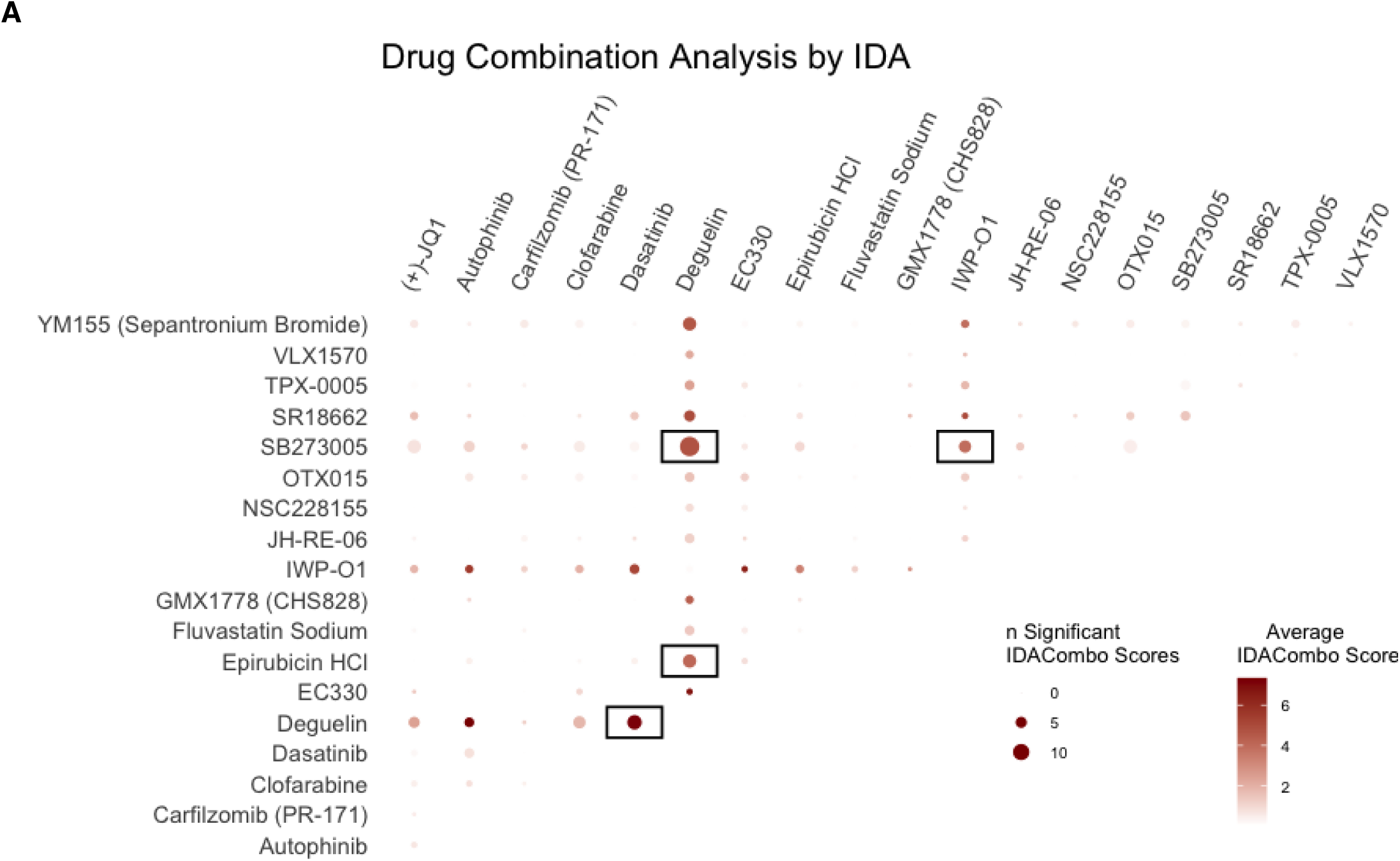
IDACombo accurately predicts combination efficacy. A. Bubble chart of the average IDACombo scores of all hypothetical drug combinations based on the follow-up panel viability data, including the drugs that validated based on the high-throughput screening data and the control drugs. Only IDACombo scores at “effective drug concentrations” were averaged, meaning any monotherapy concentration for a drug that lowered the average viability of the cells to at least 90%. Combinations were highlighted if the average IDACombo score was greater than 2 and they were significant across at least 5 dose combinations.

We sought to determine whether IDACombo’s predictions would match our experimental findings, both for effective and ineffective combinations. The combination predicted by IDACombo to be most effective was deguelin and SB273005 (Fig. 4, Supp. Fig. 12B). We have shown that this combination is indeed more effective than either drug as a single agent (Fig. 3F). Further, drug pairs that we experimentally found to be ineffective were also not identified as promising by IDACombo’s predictions. Thus, IDACombo was able to predict the effectiveness of second-line drug combinations in the context of heterogeneous populations of therapy resistant cells.

### Differential sensitivities can be lost as cells become resistant

Resistant clones arise from individual, rare primed cells in the original drug-naive population. As demonstrated earlier, differences in these primed cells are propagated and amplified to result in the differences between resistant clones^19, 20^. Thus, a natural question is whether the unique drug sensitivities apply to both the originating primed cells and the subsequent resistant clone.^19, 20, 36^. It is possible that the primed cells have the same drug sensitivities as the resistant clones, or that resistant clones either lose sensitivities that the primed cells exhibit or gain new sensitivities as the cells adapt to the primary treatment.

We first wanted to check if there were instances in which a drug sensitivity was present in the primed cells but lost during the acquisition of full therapy resistance. We have previously shown that the cellular adaptation that accompanies therapy resistance occurs within the first two weeks of treatment^20^. This critical window provided an ideal opportunity to test how these differential drug sensitivities change during adaptation. We set up a colony formation assay as follows: we first plated drug-naive cells and initiated treatment with 1 µM vemurafenib (Fig. 5A). We then created several experimental arms: (1) vemurafenib alone for the full five weeks as a control; (2) vemurafenib plus a secondary drug added simultaneously from day 0; and (3-5) vemurafenib followed by the addition of a secondary drug after one, two, or three weeks (Fig. 5A). We monitored and quantified colony formation for all conditions at three, four, and five weeks after the initial treatment. We used the drugs we selected previously based on the two distinct sensitivity patterns observed in our high-throughput screening data at the lowest dose at which we had observed differential effects among the resistant clones: drugs with specific effects (SR18662 and saracatinib) and drugs with broad moderate to strong effects (dasatinib and (+)-JQ1) across all resistant clones (Supp. Fig. 6A, Supp. Fig. 8B-D). We also included the negative control drug (S3I-201) (Supp. Fig. 8E).

**Figure 5.**
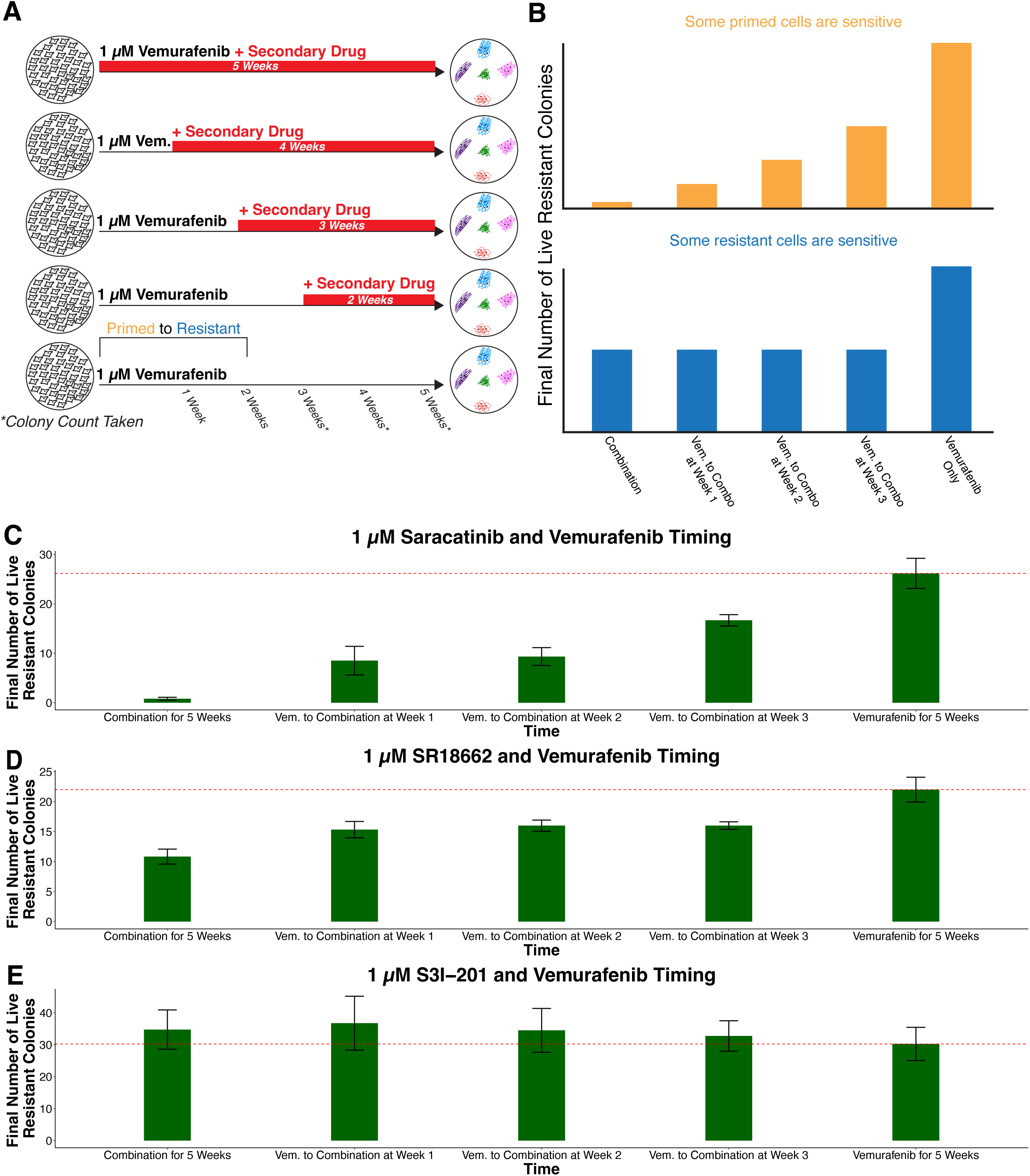
Resistant clones can lose drug sensitivities. A. Cartoon schematic showing the experimental design for the timing experiments. Briefly, drug-naive cells were plated and then treated with either vemurafenib only for five weeks, vemurafenib in combination with another drug for five weeks, or vemurafenib for one week with another drug added for the remaining four weeks, vemurafenib for two weeks with another drug added for the remaining three weeks, or vemurafenib for three weeks with another drug added for the remaining two weeks. Colony counts were quantified and compared at three, four, and five weeks for all conditions. B. Cartoon schematics showing the hypothetical number of colonies at the final week 5 time point if a secondary drug was added at each time point shown in panel C, for both the case in which some resistant cells are sensitive as well as the case in which some primed cells are sensitive. C. Bar graph of the number of live colonies at five weeks after treatment with vemurafenib only, vemurafenib and saracatinib, or vemurafenib followed by addition of saracatinib after one, two, or three weeks. The number of live colonies is the average across three experiments, each of which was done in duplicate (n = 6, n = 10 for combination treatment). D. Bar graph of the number of live colonies at five weeks after treatment with vemurafenib only, vemurafenib and SR18662, or vemurafenib followed by addition of SR18662 after one, two, or three weeks. The number of live colonies is the average across three experiments, each of which was done in duplicate (n = 6). E. Bar graph of the number of live colonies at five weeks after treatment with vemurafenib only, vemurafenib and S3I-201, or vemurafenib followed by addition of S3I-201 after one, two, or three weeks. The number of live colonies is the average across two experiments, each of which was done in duplicate (n = 4).

If a drug sensitivity across all colonies was lost over time, then we would expect to see that early co-treatment would result in fewer total resistant colonies than a later treatment; i.e., resistant cells are in general less susceptible to drugs than primed cells (Fig. 5B). We saw such an effect with saracatinib (1 μM) nearly blocking colony formation when added from the beginning, reduced colony numbers by approximately 60% when added after 7 or 14 days of vemurafenib treatment, and reduced colony numbers at 5 weeks by approximately 40% when added after 21 days (Fig. 5C). This effect was of particular note given that we used a lower dose (1 μM) than where we observed differential effects among resistant clones (10 μM) to minimize off-target effects. Similarly, the addition of dasatinib (0.1 μM) blocked colony formation when added from the beginning or after 7 days of vemurafenib treatment, with a slight increase in the number of colonies when added after 14 or 21 days of vemurafenib (Supp. Fig. 13A). Although co-treatment with vemurafenib and (+)-JQ1 (0.1 μM) did not completely block colony formation, we did observe an increase in the number of colonies at 5 weeks when (+)-JQ1 was added after 21 days of vemurafenib (Supp. Fig. 13B).

In contrast, if the number of ultimate resistant colonies was the same regardless of whether the secondary treatment was co-applied with vemurafenib or applied later, that would suggest that either the primed cells are either equally or less susceptible to the drugs than resistant cells (Fig. 5B). For example, whether SR18662 (1 μM) was added after 7, 14, or 21 days of vemurafenib treatment or combined with vemurafenib from the beginning, it resulted in approximately a 50% reduction in colony numbers at 5 weeks compared to vemurafenib-only treatment (Fig. 5D). The negative control S3I-201 (1 μM) had no effect on colony formation regardless of when it was added (Fig. 5E).

Although saracatinib and dasatinib likely have a common target, the addition of dasatinib at earlier time points resulted in fewer colonies at 5 weeks than the addition of saracatinib, suggesting that the sensitivity to saracatinib was lost more quickly^24–26^. Given that we observed a different sensitivity profile among resistant clones between saracatinib—where saracatinib strongly affected Resistant Clone 2 but not others— and dasatinib—where dasatinib strongly affected Resistant Clone 2 and had a moderate effect on others—the slower loss of sensitivity to dasatinib suggests that the other targets of dasatinib could be unrelated to the development of resistance.

Collectively, these findings demonstrated that differential drug sensitivities among resistant clones can be lost during development.

## DISCUSSION

Therapy resistance remains one of the greatest challenges in cancer treatment. Previous work has shown that even amongst a pool of clonal cells, non-genetic heterogeneity can lead to resistance as well as morphological and transcriptional variation among the resultant resistant cells. Our findings demonstrate that genetically identical therapy-resistant melanoma cells display remarkably distinct drug sensitivity profiles, with each resistant clone exhibiting a distinct fingerprint of second-line drug vulnerabilities. This heterogeneity exists despite the absence of genetic mutations driving resistance, highlighting the critical need for methods beyond whole genome sequencing to identify potential second-line inhibitors of therapy resistant cells.

Our subpopulation-directed synergy approach could substantially expand the repertoire of viable second-line therapies for patients with resistant melanoma. While our screening identified several “universal killers” effective against all resistant clones, these drugs typically targeted fundamental cellular pathways that do not offer much discrimination between cancer and normal cells. In contrast, the premise of our approach is that it may uncover specific vulnerabilities unique to resistant subpopulations that are more cancer-specific and thus less toxic to non-cancerous cells. Furthermore, by simultaneously targeting multiple resistance mechanisms, our approach may preemptively block the emergence of further resistance. Using IDACombo, we can also accurately predict the drug combinations most likely to be effective. Our findings establish the conceptual basis for the population-based synergy approach to developing second-line strategies that can draw from a much broader pool of potential therapeutic agents.

The distinct drug sensitivity patterns we observed are not confined to melanoma. Similar phenomena have been documented in non-small cell lung cancer (NSCLC), where Ramirez et al. (2016) demonstrated heterogeneity in drug sensitivity among clonal resistant cells^18^. Additionally, Goyal et al. 2023 showed that clonal resistant breast cancer cells display morphological and transcriptional variability, suggesting that non-genetic resistance heterogeneity may be a common feature across multiple cancer types^19^. Our approach of high-throughput and targeted screening of resistant clones provides a framework that could be applied to other cancers to identify second-line drug vulnerabilities otherwise masked by population averages.

Our temporal analysis revealed that differential drug sensitivities can be lost during resistance development. Previous work has shown that knocking out genes can increase or decrease the number of resistant colonies by either changing the frequency of the cells “primed” to become resistant, or enabling non-”primed” cells to develop resistance^36^. The results of these genetic alterations suggest distinct temporal windows of vulnerability that can be targeted with specific drugs. Intriguingly, some resistant clones selectively lose these pre-existing vulnerabilities, as demonstrated by saracatinib, which blocks early colony formation but shows differential effects among established resistant clones. The expression of AXL in Resistant Clone 2 correlated with its heightened sensitivity to dasatinib, consistent with previous findings by Rebecca et al. 2023^27^. Furthermore, our observation that Src inhibition affects resistant cells aligns with the finding by Torre et al. 2021 that knocking out *SRC* in a clonal, drug-naive cell population resulted in a reduction in the number of resistant colonies after treatment with vemurafenib, while knocking out *CSK* (a suppressor of SRC), increased the number of resistant colonies^36^. This temporal dynamics of drug sensitivity suggests that precise targeting of vulnerabilities that are lost over time could inform more effective treatment strategies in the clinic.

Our work reveals that considering population heterogeneity among resistant cells can uncover therapeutic opportunities masked when the resistant cells are assumed to be homogeneous. By characterizing the unique drug sensitivity profiles of genetically identical resistant clones, we identified vulnerabilities that can be exploited through combination therapies to more effectively combat resistance. This approach provides a framework for developing more precise and effective strategies to overcome therapy resistance in melanoma and potentially other cancers.

## ACKNOWLEDGEMENTS

The authors thank current and former members of the Raj lab for insightful discussions related to this work, particularly Y. Goyal, C. Triandafillou, A.E. O’Farrell, V. Ayyappan, G. Kinsler, L. Van Eyndhoven, Y. Heyman, N. Jain, and S. Reffsin; the Holzman lab at the University of Pennsylvania, particularly H. Wong, for the use of their microplate reader; the Molecular Screening & Protein Expression Facility at the Wistar Institute, particularly J. Cassel and J. Salvino, for assistance with high-throughput and targeted screening; and the Genomics Facility at the Wistar Institute, particularly S. Majumdar and S. Widura, for assistance with sequencing. Funding support for The Wistar Institute core facilities was provided by Cancer Center Support Grant P30 CA010815. This work was supported by National Institutes of Health instrument award S10OD030245-01 for the acquisition of the Echo 650 Acoustic liquid handler. G.T.B. acknowledges support from NSF GRFP DGE-2236662. R.H.B., J.L., and P.T.R acknowledge support from NIH Medical Scientist Training Program T32GM007170. R.H.B. acknowledges support from NIH Training Grant In Computational Genomics T32HG000046. R.F.G. acknowledges support from NIH Training Grant T32-CA009138-46. M.H. acknowledges support from NIH grants U54 CA224070 and P30 CA010815, as well as the Dr. Miriam and Sheldon G. Adelson Medical Research Foundation. R.S.H acknowledges support from NIH/NCI Grants R01CA204856, University of Minnesota (UMN) Office of Academic Clinical Affairs (OACA) Grant-in-Aid Program (GIA) award, NIH/NCI R01CA229618, UMN OACA Faculty Research Development grant, UMN Masonic Cancer Center CRTI Exceptional Translational Research award, and UMNCollege of Pharmacy SURRGEaward. A.R. acknowledges support from a center grant from the Mark Foundation for Cancer Research, NIH Director’s Transformative Research Award R01 GM137425, NIH R01 CA238237, NIH R01 CA232256, and NIH 4DN U01 DK127405.

## AUTHOR CONTRIBUTIONS

G.T.B. and A.R. conceived the project. A.R. and G.T.B. designed all experiments. G.T.B. performed and analyzed all experiments, supervised by A.R. G.T.B. grew cells for high-throughput and targeted screens, which were conducted by the Wistar Molecular Screening & Protein Expression Facility. R.H.B., J.L., and P.T.R. assisted G.T.B. with the manual isolation of resistant cell lines. M.J.A. and J.L. assisted G.T.B. with drug timing and drug combination experiments. R.H.B., J.L., and P.T.R. assisted G.T.B. with bulk RNA-sequencing experiments and analysis. R.F.G. predicted combination efficacies using IDACombo. G.T.B. prepared all illustrations used in this study. G.T.B. and A.R. wrote the manuscript with input from all authors.

## DATA AND CODE AVAILABILITY

All data and code are accessible at: https://www.dropbox.com/scl/fo/ypo6kxbx314nwbtwg1ei8/APgWeR0t0Y2-o5ETx89aK4g?rlkey=pa2jv5wldexz43m6t1rl27erx&st=b935hoge&dl=0

## METHODS

### Cell Culture

WM989 melanoma cells from the laboratory of Meenhard Herlyn at The Wistar Institute were twice bottlenecked to generate the WM989 A6-G3 subclone, first described in Shaffer et al. *Nature* 2020. This bottlenecking was done to minimize genetic heterogeneity. This subclone was validated by DNA short tandem repeat (STR) microsatellite fingerprinting at the Wistar Institute^37^. The cells were cultured in TU2% media containing 78% MCDB (Sigma M7403), 20% Leibovitz’s L-15 media (Life Technologies Inc., 11415064), 2% FBS, 1.68 mM CaCl2, and 50 U ml-1 penicillin and 50 μg ml-1 streptomycin (Invitrogen 15140122). All cell lines were passaged using 0.05% trypsin-EDTA (Invitrogen 25300054). Resistant clones were cultured in TU2% media containing either 1 µM vemurafenib or 250 nM dabrafenib/2.5 nM trametinib, and had their media changed the day after plating to remove residual trypsin as they could not be centrifuged to do so. All cell lines tested negative for mycoplasma. Cells were cultured in Corning Falcon plates (12-well: 08-722-29, 24-well: 08-722-1) for all secondary drug timing and combination experiments. Cells were cultured in white walled, clear bottom Corning Falcon plates (96-well: 353377) for all orthogonal high-throughput screening validation experiments.

### Drug Stocks

Stock solutions of drugs were prepared in DMSO at the specified concentrations: 50 μM trametinib (Selleckchem S2673), 1 mM dabrafenib (Cayman Chemical 16989-10), 10 mM vemurafenib (PLX4032) (Selleckchem S1267), 10 mM bortezomib (Selleckchem S1013), 10 mM epirubicin HCl (Selleckchem S1223), 10 mM S3I-201 (Selleckchem S1155), 10 mM (+)-JQ1 (Selleckchem S7110), 10 mM saracatinib (Selleckchem S1006), 10 mM birabresib (OTX-015) (Selleckchem S7360), 10 mM camptothecin (CPT) (Selleckchem S1288), 10 mM dasatinib (Selleckchem S1021), 10 mM clofarabine (Selleckchem S1218), 10 mM SB273005 (Selleckchem S7540), 10 mm SR18662 (Selleckchem S8900), 10 mM GMX1778 (Selleckchem S8117), 10 mM IWP-O1 (Selleckchem S8645), 10 mM Deguelin (Selleckchem S8132), and 10 mM Sepantronium Bromide (YM155) (Selleckchem S1130). We prepared aliquots (6-10 µL) of each drug and stored them at −20 °C to minimize freeze–thaw cycles.

### Manual Isolation of Resistant Clones

WM989 A6-G3 cells were plated at a density of 50,000 cells/plate on 15-cm tissue culture plates to minimize overlap of colonies. Cells were treated with 1 µM vemurafenib (PLX4032, Selleck Chemicals, S1267) for four weeks or 250 nM dabrafenib/2.5 nM trametinib (Cayman Chemical 16989-10/Selleckchem S2673) for six weeks to allow resistant colonies to develop and expand. Plates were scanned under a bright field tissue culture microscope to identify colonies that were sufficiently isolated from other colonies and singletons (cells which survive targeted therapy treatment but do not divide to form colonies). These distant colonies were physically isolated and dissociated via treatment with 50 µL of 0.05% trypsin for 5-10 minutes, with time varying between colonies as some took longer to dissociate than others. Colony suspensions were then plated in 96-well plates containing 150 µL of Tu2% medium with either 1 µM vemurafenib or 250 nM dabrafenib/2.5 nM trametinib added, as appropriate.

After cells had settled the next day, 150 µL of media was replaced to remove residual trypsin without disrupting the cells. Isolated resistant colonies were monitored each day to determine health and confluence, with the media being changed every 3-5 days. Treatment with 1 µM vemurafenib or 250 nM dabrafenib/2.5 nM trametinib was maintained throughout the entire isolation and expansion process. As colonies reached 70-80% confluence, they were serially expanded into 24-well, 12-well, 6-well, 6-cm, and then 10-cm plates. The resulting resistant cell lines, dubbed resistant clones, enabled further analysis.

### Bulk RNA Sequencing and Analysis of Resistant Clones

The first bulk sequencing run was conducted as described in Goyal et al. 2023. The second bulk RNA sequencing run was conducted as follows. Cells were collected at a density of 75,000 per sample as resistant clones were expanded, and stored in QIAzol until all samples had been collected. We then conducted standard bulk paired-end (37:8:8:38) RNA-seq using RNeasy Micro (Qiagen 74004) for RNA extraction, NEBNext Poly(A) mRNA Magnetic Isolation Module (NEB E7490L), NEBNext Ultra II RNA Library Prep Kit for Illumina (NEB E7770L), NEBNext Multiplex Oligos for Illumina (Dual Index Primers Set 1) oligonucleotides (NEB E7600S), and an Illumina NextSeq 1000/2000 P4 XLEAP-SBS 100 cycles kit (Illumina 20100994), as previously described^22, 38^. We sequenced each sample at a depth of approximately 50 million reads on a NextSeq2000 with the help of the Wistar Institute Genomics Facility. Prior to extraction and library preparation, the samples were randomized to avoid any experimental and human biases. For both runs, we pseudoaligned RNA-seq reads to the human genome (hg19) with Kallisto to obtain transcripts per million. Data was exported to R for downstream analysis.

### High-Throughput Screening of Resistant Clones

Resistant Clones 1, 2, 3, and 4 were used for high-throughput screening at The Wistar Institute’s Molecular Screening & Protein Expression Facility. We used a pan-cancer library containing 2240 drugs and small molecules with known pathways and targets against different cancer types (Supp. Table 1). We used a four-point dose curve of 0.01 µM, 0.1 µM, 1 µM, and 10 µM for each drug. An Echo 650 Acoustic liquid handler was used to transfer the library.

The screen was conducted by plating cells at a density of 1000 cells/well in 384-well plates and incubating with the library for seven days. The readout was cell viability as measured by CellTiter-Glo with a BMG ClarioStar Plus plate reader, with the luminescence readout normalized to a positive control readout (cells treated with the protease inhibitor Bortezomib or Epirubicin HCl) and a negative control readout (cells treated with the vehicle DMSO) (Fig. 1B).

We performed the screen in two batches: Resistant Clones 1 and 2 in Screen A, and Resistant Clones 3 and 4 in Screen B, with the drug-naive WM989 A6-G3 cells in both (Fig. 1B). Note that in Screen A, we performed the screen for resistant and drug-naive cell lines both with and without vemurafenib, while in Screen B the resistant clones were screened together with vemurafenib and the drug-naive cells were screened both with and without vemurafenib (Fig. 1B). (We did not screen the resistant clones without vemurafenib in Screen B as analysis of Screen A data showed that the viability values of the library with and without vemurafenib were highly correlated, suggesting these conditions provided largely redundant information (Supp. Fig. 2B)).

### Validation of High-Throughput Screening

We selected a set of eight drugs for validation of the high-throughput screening results: epirubicin HCl, S3I-201, clofarabine, dasatinib, camptothecin, saracatinib, OTX-015, and (+)-JQ1. We used a four-point dose curve of 0.01 µM, 0.1 µM, 1 µM, and 10 µM for each drug. The experiments were conducted by plating cells at a density of 5000 cells/well in 96-well plates. The plates were imaged on an Incucyte S3 Live Cell Imaging Analysis System (Sartorius) with a 4x objective and incubation with each drug was begun 12-16 hours after the cells had been plated. Cells were incubated with the library for seven days and the media was changed once during the incubation period, after 3-4 days. After seven days, the plates were imaged again on an Incucyte S3 and the CellTiter-Glo assay (Fig. 1B) was used to measure cell viability. A DTX880 Multimode Detector was used to measure the luminescence signal generated by the CellTiter-Glo assay.

### Calculation of Significance Thresholds

We established statistical thresholds for the amount of variability in viability to expect by chance by using the drug sensitivities of drug-naive cells, which were included in both Screen A and B, as an across-batch test of differences arising from technical variability. For each drug, we measured the difference in viability between these replicates. Measuring the variance in this difference across all drugs at doses of 0.01 µM, 0.1 µM, and 1 µM, we adopted two standard deviations (representing an approximately 20 percentage point difference in viability) as our threshold for significance, which corresponded to an approximately 95% confidence interval. We also used Screen A’s within-batch replicates (Resistant Clones 1 and 2 with and without vemurafenib) as another test of technical variability that did not reflect the batch differences. Again, for each drug we measure the difference in viability between these replicates. Measuring the variance in this difference across all drugs at each dose, we found that two standard deviations also represented an approximately 20 percentage point difference in viability (corresponding to an approximately 95% confidence interval) at doses of 0.01 µM, 0.1 µM, and 1 µM. However, we found that the difference in viability representing two standard deviations was much higher in the across-batch test versus the within-batch test. Therefore, we excluded all drug-dose conditions at a dose of 10 µM from further analysis.

### Distribution of Pathways in Top Hits of the High-Throughput Screening Data

We first found all drug-dose conditions to which at least one individual cell line was highly sensitive (viability < 30% in at least one line), then filtered out all drug-dose conditions that to which all cells were sensitive (viability < 50% across all cell lines). We chose a threshold of 50% as it passed the true difference threshold of 20% identified in our cross-batch analysis, so all remaining drug-dose conditions under 30% would satisfy this threshold.

For each resistant clone and the drug-naive cells, we calculated how often the pathway being inhibited by each drug was represented in the subset of drug-dose conditions and normalized this count to the total number of drugs in the panel targeting that pathway. We then ranked the top eight pathways for each of the resistant clones and the drug-naive cells.

### Gene Set Enrichment Analysis of Metaprograms

Using the fgsea package in R, we conducted GSEA on the bulk RNA sequencing data of the resistant clones we used in high-throughput screening, as well as the drug-naive cells and additional resistant clones representing diverse resistant types^19, 21, 39^. We used the metaprograms described in Boe et al. 2024 as the gene sets. We plotted the normalized enrichment scores in a heatmap. We selected a set of resistant clones for high-throughput screening representing expression of diverse metaprograms.

### Comparison of Transcriptomic and Drug Sensitivity Distances

To quantify the correlation between drug sensitivity and transcriptome, we calculated distances between each pair of resistant clones in both transcriptomic and drug sensitivity PCA space. We first conducted PCA on bulk RNA sequencing data of the resistant clones we used in high-throughput screening, as well as the drug-naive cells and additional resistant clones representing diverse resistant types^19, 21^. We then calculated the Euclidean distance in PCA space between each pair of Resistant Clones 1 - 4.

We next conducted PCA on the high-throughput screening data for each drug-dose condition (excluding those at doses of 10 µM) for the Resistant Clones 1 - 4 and the drug-naive cells. We calculated the Euclidean distance in PCA space between each pair of Resistant Clones 1 - 4. We then compared these distances to the distance calculated in transcriptomic space, and calculated both the Pearson’s and Spearman’s correlation coefficients, using a p-value of 0.05 as a cutoff for a statistically significant correlation.

### Comparison of Resistance Program and Sensitivity Profiles

We wanted to know how closely associated the resistance programs were with the sensitivity profiles of the resistant clones. We calculated the number of differential drug-dose conditions between each pair of resistant clones that passed our confidence threshold (i.e. a difference of more than 20 percentage points in viability), and compared these numbers to the number we would expect to see pass the confidence threshold when comparing the drug-naive cells in Screen A and B, as well as when comparing Resistant Clones 1 and 2 with and without vemurafenib in Screen A.

### Distribution of Type of Killer in the High-Throughput Screening Data

We categorized all remaining drug-dose conditions at 1 µM, 0.1 µM, or 0.01 µM (6720 total) as universal, shared, or unique killers. We defined universal killers as killing all clones, shared killers as killing multiple clones and not killing at least one other clone, and unique killers as killing only one clone and not killing at least one other clone. To maximize the degree of specificity, we used a threshold of < 30% viability for a drug-dose condition to kill a clone and > 70% to leave a clone alive.

### Targeted Screening of Additional Resistant Clones

We selected a set of 24 drugs for targeted screening with additional resistant clones. We included the original four clones that were used in the high-throughput screens, as well as five additional vemurafenib-resistant clones and four additional dabrafenib/trametinib-resistant clones. We also included the drug-naive WM989 A6-G3 cells. The screen was conducted by The Wistar Institute’s Molecular Screening & Protein Expression facility. Resistant clones were screened in media containing either vemurafenib or dabrafenib/trametinib, depending on their primary resistance. Drug-naive cells were screened without any additional inhibitor (only the vehicle DMSO). An Echo 650 Acoustic liquid handler was used to transfer the library.

The follow-up panel was conducted in one phase. Cells were plated at a density of 1000 cells/well in 384-well plates and incubated with the panel for seven days. The CellTiter-Glo Assay was then used to measure cell viability using a BMG ClarioStar Plus plate reader, and the readout was normalized to a positive control value (epirubicin HCl) and a negative control value (the vehicle DMSO). Two technical replicates were used for this panel. An 8-point logarithmic dose curve was used, with the starting dose varying based on the dose at which we had seen differences among the resistant lines in the high-throughput screens.

### Euclidean Distance Matrix Calculation

To quantify the differences in the sensitivity profiles of the resistant clones in Follow-Up Panel A, we calculated the pairwise Euclidean distance, using the 16 drugs at the doses of interest in Follow-Up Panel A that validated based on the high-throughput screening data, as well as dasatinib at a dose of 0.1 µM, for each pair of resistant clones (13 clones total). Although dasatinib was initially included in this panel as a control drug, we included it in this analysis due to the observation that Resistant Clone 2, which was uniquely sensitive to dasatinib in the high-throughput screening data, highly expresses the canonical resistance marker *AXL*, which has been linked to sensitivity to dasatinib^23, 27, 28^.

### Secondary Drug Timing

For all timing experiments, cells were plated at a density of 10,000 cells per well in either 12-well or 24-well plates. The plates were imaged on an Incucyte S3 Live Cell Imaging Analysis System (Sartorius) with a 4× objective and drug treatment was begun 12-16 hours after the cells had been plated. The experimental arms were as follows: (1) vemurafenib alone for the full five weeks as a control; (2) vemurafenib plus a second drug added simultaneously from day 0; and (3-5) vemurafenib followed by the addition of a second drug after one, two, or three weeks. A total concentration of 0.1% DMSO was maintained across all experimental arms. Media was changed every 3-4 days, and plates were imaged once a week on the Incucyte S3.

### Hypothetical Drug Combination Analysis

We selected a subset of 10 drugs from those in Follow-Up Panel A that validated based on the high-throughput screening data and killed some resistant clones and not others, rather than drugs that had similar effects (or lack thereof) across all resistant clones. While we used a stringent 30% viability threshold elsewhere in our analyses to identify high-confidence single-drug effects, for this combinatorial approach we employed a more inclusive 50% viability threshold for a clone to be considered killed by a drug. This lower threshold was selected to capture both highly potent (< 30%) as well as moderate (30 - 50%) drug effects, allowing us to identify synergistic drug pairs where one drug may have moderate activity against certain clones that complement the strong activity of another drug against different clones. After assigning each clone as killed or not killed for each drug, we totaled the number of resistant clones killed by each drug and calculated the theoretical total of resistant clones killed by each combination of drugs, assuming independent effects with no double-counting of clones affected by both drugs. We then selected a subset of combinations which hypothetically killed all or all but 1-2 resistant clones for validation.

### Secondary Drug Combinations

For all combination experiments, cells were plated at a density of 10,000 cells per well in 12-well plates. The plates were imaged on an Incucyte S3 Live Cell Imaging Analysis System (Sartorius) with a 4× objective and drug treatment was begun 12-16 hours after the cells had been plated. The experimental arms were as follows: (1) primary drug (vemurafenib or dabrafenib/trametinib) for the full six weeks as a control; (2) primary drug (vemurafenib or dabrafenib/trametinib) followed by the addition of a secondary drug after four weeks; (3) primary drug (vemurafenib or dabrafenib/trametinib) followed by the addition of another secondary drug after four weeks; and (4) primary drug (vemurafenib or dabrafenib/trametinib) followed by the addition of both secondary drugs after four weeks. A total concentration of 0.1% DMSO was maintained across all experimental arms. Media was changed every 3-4 days, and plates were imaged once a week on the Incucyte S3.

### Image Analysis

Image analysis for all drug timing and combination experiments was conducted using NimbusImage (https://github.com/Kitware/UPennContrast, https://www.nimbusimage.com/). For the drug combination experiments, colonies were segmented using the SegmentAnything worker at the four week time point (https://github.com/facebookresearch/segment-anything).

The Propagate worker was then used to automatically segment colonies at the five and six week time points using SegmentAnything and the four week time point annotations. These propagated annotations were manually reviewed for accuracy. For the drug timing experiments, colonies were segmented using the SegmentAnything worker at the three week time point. The Propagate worker was then used to automatically segment colonies at the four and five week timepoints. Colonies were manually categorized as live or dead based on morphology and the total colony counts at each time point were computed.

### Independent Drug Action (IDA) Analysis and IDACombo

The IDA analysis and generation of IDACombo Scores was performed using the pipeline developed in Ling et al, 2020. (https://pubmed.ncbi.nlm.nih.gov/33203866/).

Briefly, IDA assumes the effect of a combination on a single sample is equal to the effect of the most efficacious drug in this combination (i.e. it assumes no additivity or synergy in any single sample), but that across samples the combinations can be more effective since the sample has multiple chances of responding to a drug. We can then predict combination efficacy by IDA by selecting the maximal response the cell line has to 2 drugs. This is averaged over all cell lines to create a mean effect the drug has on a population given IDA, which can then be compared to the mean effect of each monotherapy alone to generate an IDACombo score.

IDACombo scores were generated on the drug response data for all resistant cell lines using the IDACombo R package available at https://github.com/Alexander-Ling/IDACombo/tree/master. Statistical analysis is performed by using the variance in the drug response data and performing 10,000 Monte Carlo simulations of the combination effect. This results in an average IDACombo score with 95% confidence intervals for each drug combination at every possible dose pair. To ensure the drug combination is robust, we averaged the IDACombo score over the available dose range and compared these averaged scores across all drug combinations. Combinations were highlighted if they were in the top 10% of average IDACombo scores and were significant across at least 8 dose combinations. [These values are arbitrary, but do represent the drug combinations that had both a magnitude and significance in the top 10% of all drug combinations].

[Averaging IDACombo scores over all the possible drug combinations ensures the IDACombo effect is robust and seen at multiple doses. However, since the number of doses a drug has has an impact on viability changes for every drug, this would also influence the average IDACombo scores. This is not necessarily a bad thing, since we also want to prioritize drugs that are effective at that dose range. However, we performed another analysis to ensure the best candidate drug combinations were still the best when correcting for this. Here, only IDACombo scores at “effective drug concentrations” were averaged, with “effective drug concentrations” meaning any monotherapy concentration for a drug that lowered the average viability of the cells to at least 90%.]

[Cutoffs for a dotplot figure: n_sig_IDACombo >= 8, Avg_IDACombo > 1, for “effective concentration plot it is: .n_sig_IDACombo > 5, Avg_IDACombo > 2, and ratio_sig_IDACombo > 0.5. This ratio is the ratio of the number of significant IDACombo scores [n_sig_IDACombo] to the total number of effective dose pairs for that drug combination]

## DECLARATION OF GENERATIVE AI AND AI-ASSISTED TECHNOLOGIES

During the preparation of this work, the authors used ChatGPT and Claude to generate/comment code for analyses. After using this tool or service, the authors reviewed and edited the content and take full responsibility for the content of the publication.

## REFERENCES

1. Gonzalez, D. et al. BRAF mutation testing algorithm for vemurafenib treatment in melanoma: recommendations from an expert panel: Recommendations forBRAFtesting for vemurafenib. Br. J. Dermatol. 168, 700–707 (2013).

2. Davies, H. et al. Mutations of the BRAF gene in human cancer. Nature 417, 949–954 (2002).

3. Curtin, J. A. et al. Distinct sets of genetic alterations in melanoma. N. Engl. J. Med. 353, 2135–2147 (2005).

4. Long, G. V. et al. Overall survival and durable responses in patients with BRAF V600-mutant metastatic melanoma receiving dabrafenib combined with trametinib. J. Clin. Oncol. 34, 871–878 (2016).

5. Bollag, G. et al. Clinical efficacy of a RAF inhibitor needs broad target blockade in BRAF-mutant melanoma. Nature 467, 596–599 (2010).

6. Trunzer, K. et al. Pharmacodynamic effects and mechanisms of resistance to vemurafenib in patients with metastatic melanoma. J. Clin. Oncol. 31, 1767–1774 (2013).

7. Garraway, L. A. & Jänne, P. A. Circumventing cancer drug resistance in the era of personalized medicine. Cancer Discov. 2, 214–226 (2012).

8. Nazarian, R. et al. Melanomas acquire resistance to B-RAF(V600E) inhibition by RTK or N-RAS upregulation. Nature 468, 973–977 (2010).

9. Flaherty, K. T. et al. Inhibition of mutated, activated BRAF in metastatic melanoma. N. Engl. J. Med. 363, 809–819 (2010).

10. Haas, L. et al. Acquired resistance to anti-MAPK targeted therapy confers an immune-evasive tumor microenvironment and cross-resistance to immunotherapy in melanoma. Nat Cancer 2, 693–708 (2021).

11. Atkins, M. B. et al. Combination Dabrafenib and Trametinib Versus Combination Nivolumab and Ipilimumab for Patients With Advanced BRAF-Mutant Melanoma: The DREAMseq Trial-ECOG-ACRIN EA6134. J. Clin. Oncol. 41, 186–197 (2023).

12. Pozniak, J. et al. A TCF4-dependent gene regulatory network confers resistance to immunotherapy in melanoma. Cell 187, 166–183.e25 (2024).

13. Tétu, P. et al. Benefit of the nivolumab and ipilimumab combination in pretreated advanced melanoma. Eur. J. Cancer 93, 147–149 (2018).

14. Mason, R. et al. Combined ipilimumab and nivolumab first-line and after BRAF-targeted therapy in advanced melanoma. Pigment Cell Melanoma Res. 33, 358–365 (2020).

15. Ackerman, A. et al. Outcomes of patients with metastatic melanoma treated with immunotherapy prior to or after BRAF inhibitors: Sequencing of Melanoma Therapies. Cancer 120, 1695–1701 (2014).

16. Johnson, D. B. et al. Sequencing treatment in BRAFV600 mutant melanoma: Anti-PD-1 before and after BRAF inhibition. J. Immunother. 40, 31–35 (2017).

17. Long, G. V. et al. Neoadjuvant pembrolizumab, dabrafenib and trametinib in BRAFV600-mutant resectable melanoma: the randomized phase 2 NeoTrio trial. Nat. Med. 30, 2540–2548 (2024).

18. Ramirez, M. et al. Diverse drug-resistance mechanisms can emerge from drug-tolerant cancer persister cells. Nat. Commun. 7, 10690 (2016).

19. Goyal, Y. et al. Diverse clonal fates emerge upon drug treatment of homogeneous cancer cells. Nature 620, 651–659 (2023).

20. Li, J. et al. AP-1 mediates cellular adaptation and memory formation during therapy resistance. bioRxivorg 2024.07.25.604999 (2024) doi:10.1101/2024.07.25.604999.

21. Boe, R. H., Triandafillou, C. G., Lazcano, R., Wargo, J. A. & Raj, A. Spatial transcriptomics reveals influence of microenvironment on intrinsic fates in melanoma therapy resistance. bioRxivorg 2024.06.30.601416 (2024).

22. Shaffer, S. M. et al. Rare cell variability and drug-induced reprogramming as a mode of cancer drug resistance. Nature 546, 431–435 (2017).

23. Müller, J. et al. Low MITF/AXL ratio predicts early resistance to multiple targeted drugs in melanoma. Nat. Commun. 5, 5712 (2014).

24. Warmuth, M. et al. Dual-specific Src and Abl kinase inhibitors, PP1 and CGP76030, inhibit growth and survival of cells expressing imatinib mesylate-resistant Bcr-Abl kinases. Blood 101, 664–672 (2003).

25. Skoko, J., Rožanc, J., Charles, E. M., Alexopoulos, L. G. & Rehm, M. Post-treatment de-phosphorylation of p53 correlates with dasatinib responsiveness in malignant melanoma. BMC Cell Biol. 19, 28 (2018).

26. Roskoski, R., Jr. Src protein-tyrosine kinase structure, mechanism, and small molecule inhibitors. Pharmacol. Res. 94, 9–25 (2015).

27 . Rebecca, V. W. et al. Dasatinib Resensitizes MAPK Inhibitor Efficacy in Standard-of-Care Relapsed Melanomas. bioRxiv (2023) doi:10.1101/2023.01.20.524923.

28. Flower, C. T. et al. Signaling and transcriptional dynamics underlying early adaptation to oncogenic BRAF inhibition. Cell Syst. 0, 101239 (2025).

29. Oksanen, J. et al. vegan: Community Ecology Package. Preprint at https://CRAN.R-project.org/package=vegan (2022).

30. Xu, S. et al. Use ggbreak to effectively utilize plotting space to deal with large datasets and outliers. Front. Genet. 12, 774846 (2021).

31. Xia, Y. et al. A Web Application for Predicting Drug Combination Efficacy Using Monotherapy Data and IDACombo. Journal of Cancer Science and Clinical Therapeutics 7, 253–258 (2023).

32. Ling, A. & Huang, R. S. Computationally predicting clinical drug combination efficacy with cancer cell line screens and independent drug action. Nat. Commun. 11, 5848 (2020).

33. Plana, D., Palmer, A. C. & Sorger, P. K. Independent drug action in combination therapy: Implications for precision oncology. Cancer Discov. 12, 606–624 (2022).

34. Palmer, A. C. & Sorger, P. K. Combination cancer therapy can confer benefit via patient-to-patient variability without drug additivity or synergy. Cell 171, 1678–1691.e13 (2017).

35. Foucquier, J. & Guedj, M. Analysis of drug combinations: current methodological landscape. Pharmacol. Res. Perspect. 3, e00149 (2015).

36. Torre, E. A. et al. Genetic screening for single-cell variability modulators driving therapy resistance. Nat. Genet. 53, 76–85 (2021).

37. Shaffer, S. M. et al. Memory Sequencing Reveals Heritable Single-Cell Gene Expression Programs Associated with Distinct Cellular Behaviors. Cell 182, 947–959.e17 (2020).

38. Mellis, I. A. et al. Responsiveness to perturbations is a hallmark of transcription factors that maintain cell identity in vitro. Cell Syst 12, 885–899.e8 (2021).

39. Korotkevich, G. et al. Fast gene set enrichment analysis. bioRxiv 060012 (2016) doi:10.1101/060012.

