## Supplemental Figures and Captions for "Heterogeneous therapy-resistant cancer cells have distinct and exploitable drug sensitivity profiles"

### Supplemental Figure 1

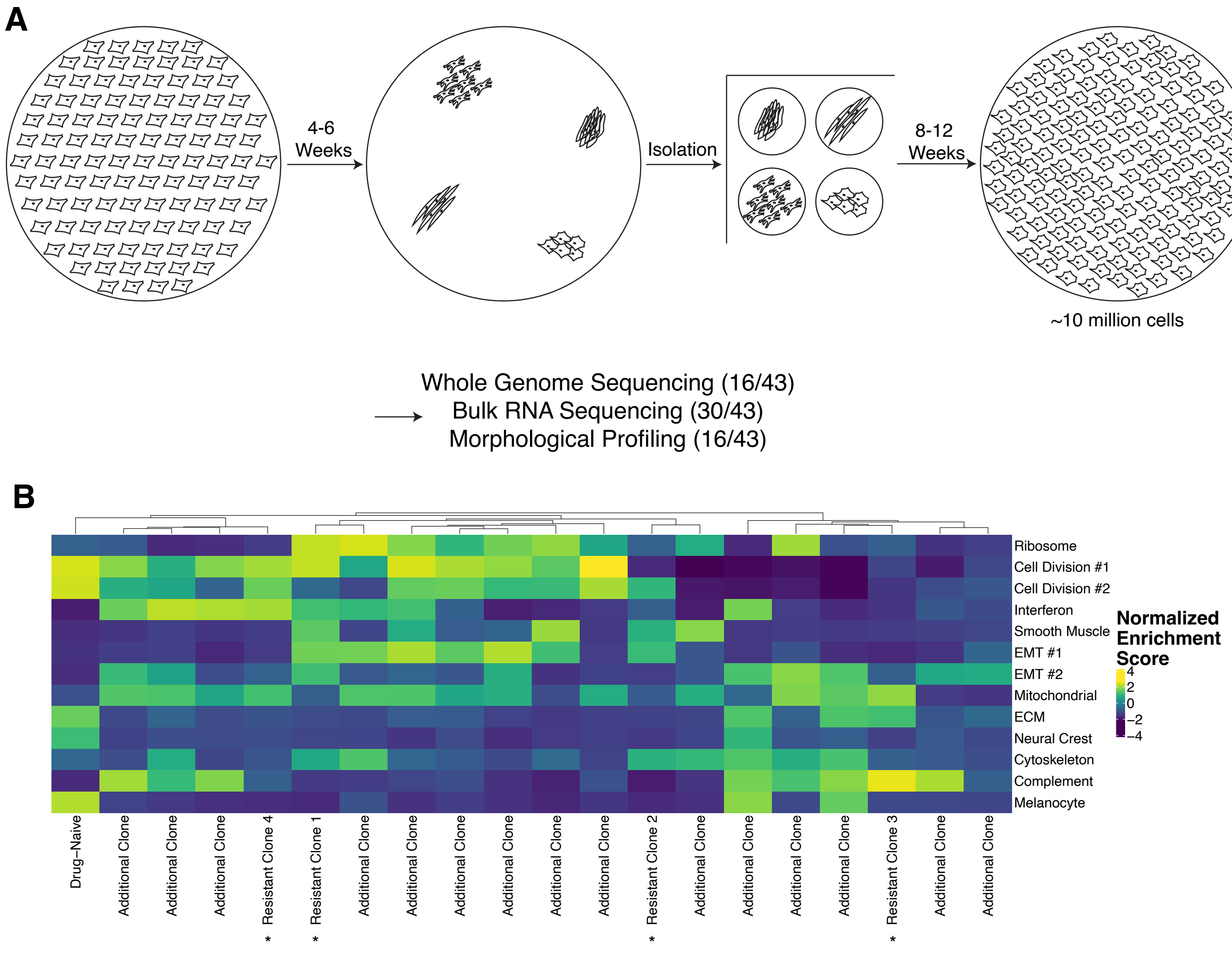


*Used in High-Throughput Screens

**A.** Cartoon schematic of the resistant clone isolation and expansion procedure. Drug-naive cells were plated sparsely and treated with targeted therapy. Once colonies had reached a sufficient size, they were manually isolated and extracted into 96-well plates. From there, they were expanded into successively larger well plates until they had grown enough to be frozen and banked. Clones were imaged to observe the morphology. Whole genome and bulk RNA sequencing were conducted on subsets of the clones.

**B.** Heatmap of the normalized enrichment scores (NES) generated by gene set enrichment analysis (GSEA) of the resistance metaprograms described in Boe et al. 2024 on a set of the resistant clones. The heatmap is clustered by cell line. Resistant Clones 1 - 4, used in the high-throughput screens, are labelled on the heatmap while the rest are labelled as Additional Clone.

**Supplemental Figure 2.**

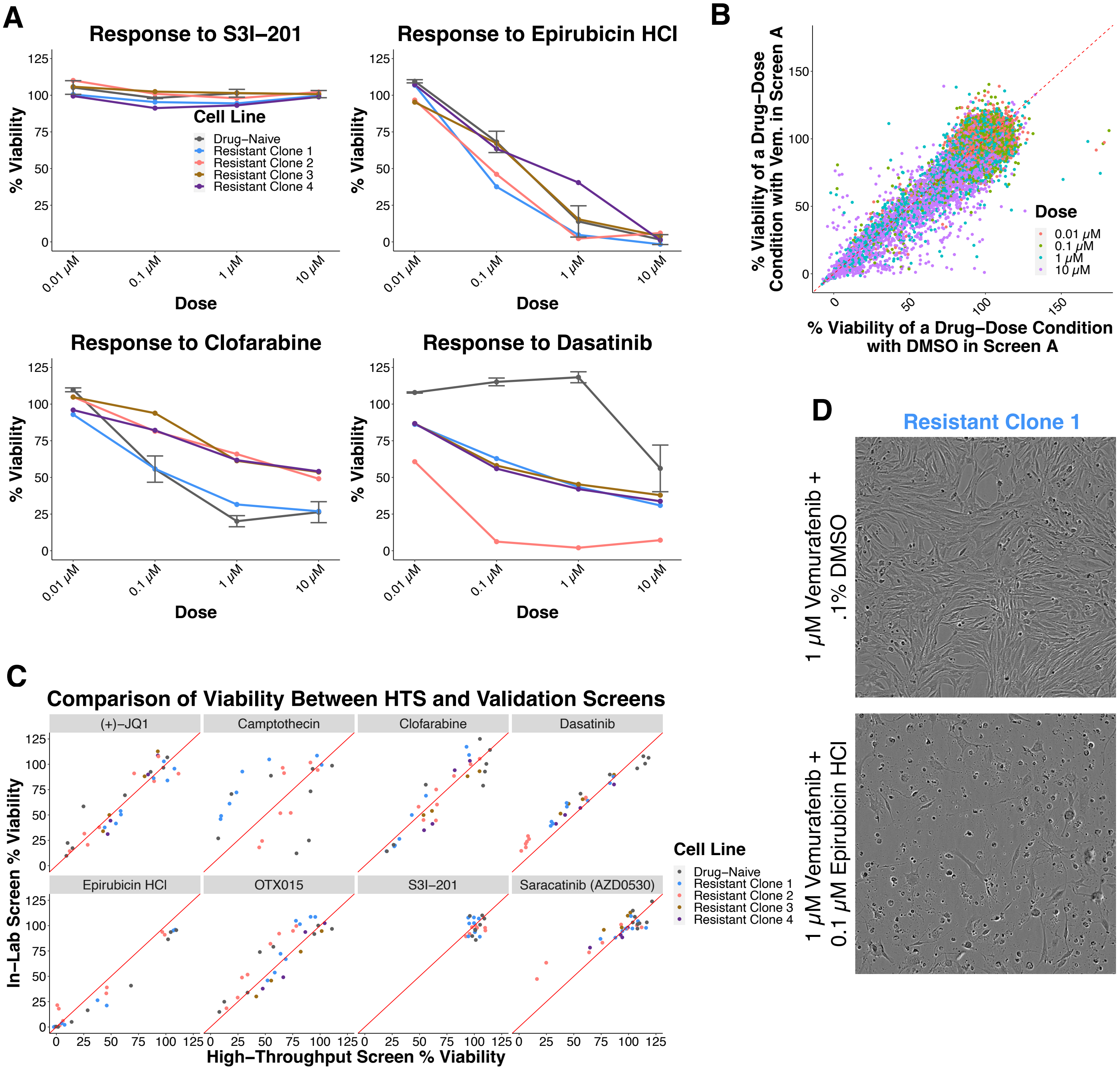

**Supplemental Figure 2.**

**A.** ***(Top Left)*** Dose curve of the response to S3I-201 for the drug-naive cells and Resistant Clones 1 - 4 in the high-throughput screens. The values shown for the resistant clones are the percent viability of the drug in combination with vemurafenib. The points for the drug-naive cells are the percent viability of the drug in combination with DMSO, and are the average of the value in each screen. ***(Top Right)*** Dose curve of the response to Epirubicin HCl for the drug-naive cells and Resistant Clones 1 - 4 in the high-throughput screens. The values shown for the resistant clones are the percent viability of the drug in combination with vemurafenib. The points for the drug-naive cells are the percent viability of the drug in combination with DMSO, and are the average of the value in each screen. ***(Bottom Left)*** Dose curve of the response to Clofarabine for the drug-naive cells and Resistant Clones 1 - 4 in the high-throughput screens. The values shown for the resistant clones are the percent viability of the drug in combination with vemurafenib. The points for the drug-naive cells are the percent viability of the drug in combination with DMSO, and are the average of the value in each screen. ***(Bottom Right)*** Dose curve of the response to Dasatinib for the drug-naive cells and Resistant Clones 1 - 4 in the high-throughput screens. The values shown for the resistant clones are the percent viability of the drug in combination with vemurafenib. The points for the drug-naive cells are the percent viability of the drug in combination with DMSO, and are the average of the value in each screen.
**B.** Scatterplot of the viabilities of drug-dose conditions on Resistant Clones 1 and 2 in Screen A in combination with vemurafenib (y axis) versus in combination with DMSO (x axis). Points are colored by dose.
**C.** Scatterplots of the eight drugs tested to validate the high-throughput screening results, faceted by drug. Each point represents a particular dose of the drug either in combination with vemurafenib or DMSO for a specific cell line, and is the average of all replicates. Data with either vemurafenib or DMSO is shown for the drug-naive cells as well as Resistant Clones 1 and 2. Data with vemurafenib only is shown for Resistant Clones 3 and 4, as these clones were not tested with DMSO in the high-throughput screens. Points are colored by cell line. The x axis is the viability in the high-throughput screens, and the y axis is the viability in the validation experiments. Each experiment was done in duplicate, and the total number of replicates for each cell line are as follows: (+)-JQ1 (Parental n = 4, Resistant Clone 1 n = 2, Resistant Clone 2 n = 4, Resistant Clone 3 n = 2, Resistant Clone 4 n = 4), Camptothecin (Parental n = 2, Resistant Clone 1 n = 2, Resistant Clone 2 n = 2), Clofarabine (Parental n = 4, Resistant Clone 1 n = 2, Resistant Clone 2 n = 4, Resistant Clone 3 n = 2, Resistant Clone 4 n = 4), Dasatinib (Parental n = 4, Resistant Clone 1 n = 2, Resistant Clone 2 n = 4, Resistant Clone 3 n = 2, Resistant Clone 4 n = 4), Epirubicin HCl (Parental n = 2, Resistant Clone 1 n = 2, Resistant Clone 2 n = 2), OTX-015 (Parental n = 4, Resistant Clone 1 n = 2, Resistant Clone 2 n = 4, Resistant Clone 3 n = 2, Resistant Clone 4 n = 4), S3I-201 (Parental n = 2, Resistant Clone 1 n = 2, Resistant Clone 2 n = 2), and Saracatinib (AZD0530) (Parental n = 4, Resistant Clone 1 n = 2, Resistant Clone 2 n = 4, Resistant Clone 3 n = 2, Resistant Clone 4 n = 4).
**D.** ***(Top)*** Brightfield image of Resistant Clone 1 treated with 1 $\mu$M vemurafenib + .1% DMSO for seven days. ***(Bottom)*** Brightfield image of Resistant Clone 1 treated with 1 $\mu$M vemurafenib + 0.1 $\mu$M Epirubicin HCl for seven days.

### Supplemental Figure 3.

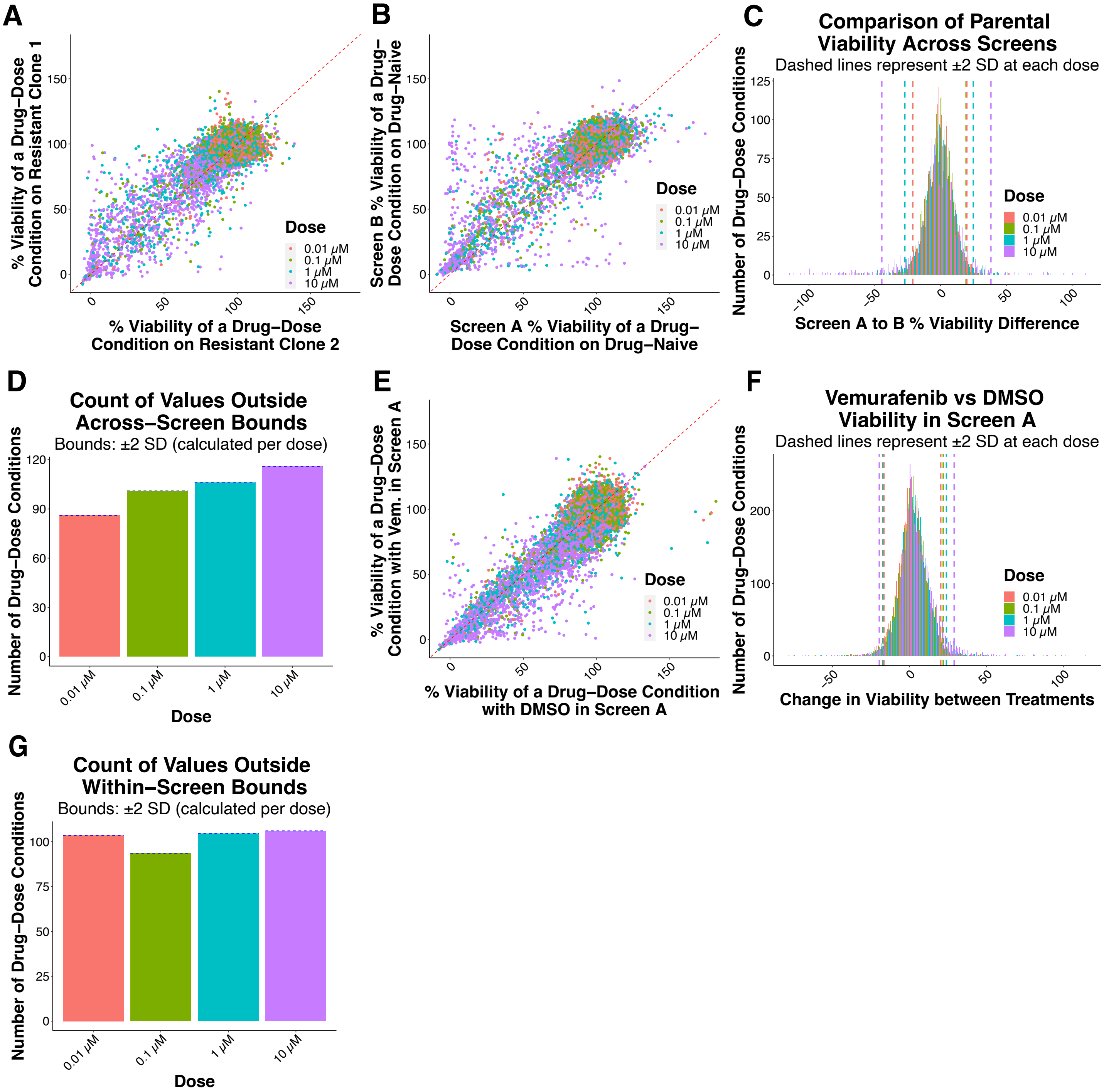

**Supplemental Figure 3.**

**A.** Scatterplot of the viabilities of drug-dose conditions on Resistant Clone 1 in Screen A in combination with vemurafenib (y axis) versus Resistant Clone 2 in Screen A in combination with vemurafenib (x axis). Points are colored by dose.
**B.** Scatterplot of the viabilities of drug-dose conditions on drug-naive cells in Screen B in combination with DMSO (y axis) versus in Screen A in combination with DMSO (x axis). Points are colored by dose. This comparison was used to estimate variation across screens.
**C.** Histogram of the difference in % viability of all drug-dose conditions on the drug-naive cells from Screen A to B. The dashed lines on the graph represent where two standard deviations falls on either side of the mean at each dose. Lines and bars are colored by dose.
**D.** Bar graph of the number of drug-dose conditions that fall outside of the bounds at two standard deviations for each dose as shown in panel C.
**E.** Scatterplot of the viabilities of drug-dose conditions on Resistant Clones 1 and 2 in Screen A in combination with vemurafenib (y axis) versus Resistant Clones 1 and 2 in Screen A in combination with DMSO (x axis). Points are colored by dose. This comparison was used to estimate variation within a screen.
**F.** Histogram of the difference in % viability between all drug-dose conditions on Resistant Clones 1 and 2 in combination with vemurafenib and in combination with DMSO . The dashed lines on the graph represent where two standard deviations falls on either side of the mean at each dose. Lines and bars are colored by dose.
**G.** Bar graph of the number of drug-dose conditions that fall outside of the bounds at two standard deviations for each dose as shown in panel F.

**Supplemental Figure 4.**

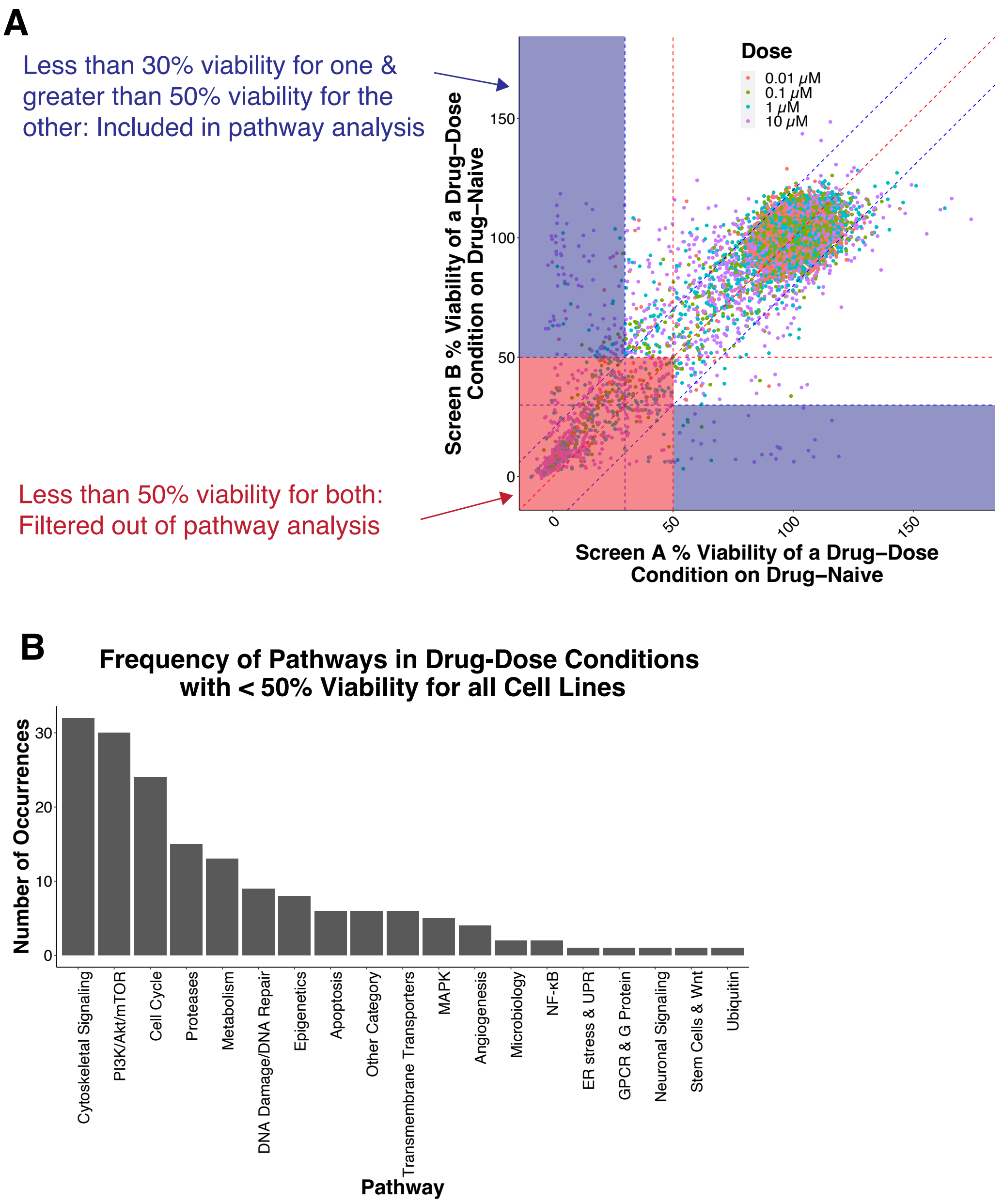


**A.** Schematic explaining why drug-dose conditions with less than 50% viability for all cell lines were filtered out. The confidence thresholds indicate that a difference of more than 20 percentage points in viability between two cell lines is likely to represent a real biological difference. Therefore, filtering out conditions with less than 50% viability for all cell lines (as indicated by the red shaded region) means that any remaining drug-dose conditions with less than 30% viability for a particular cell line (as indicated by the blue shaded region) will have a difference of more than 20 percentage points in viability as compared to at least one other cell line.
**B.** Bar plot of the frequency of the pathways among the drug-dose conditions filtered out of the rank-order plots. The 167 drug-dose conditions here are at doses of 1, 0.1, or 0.01 $\mu$M and had less than 50% viability for the drug-naive cells as well as Resistant Clones 1 - 4.

### Supplemental Figure 5.

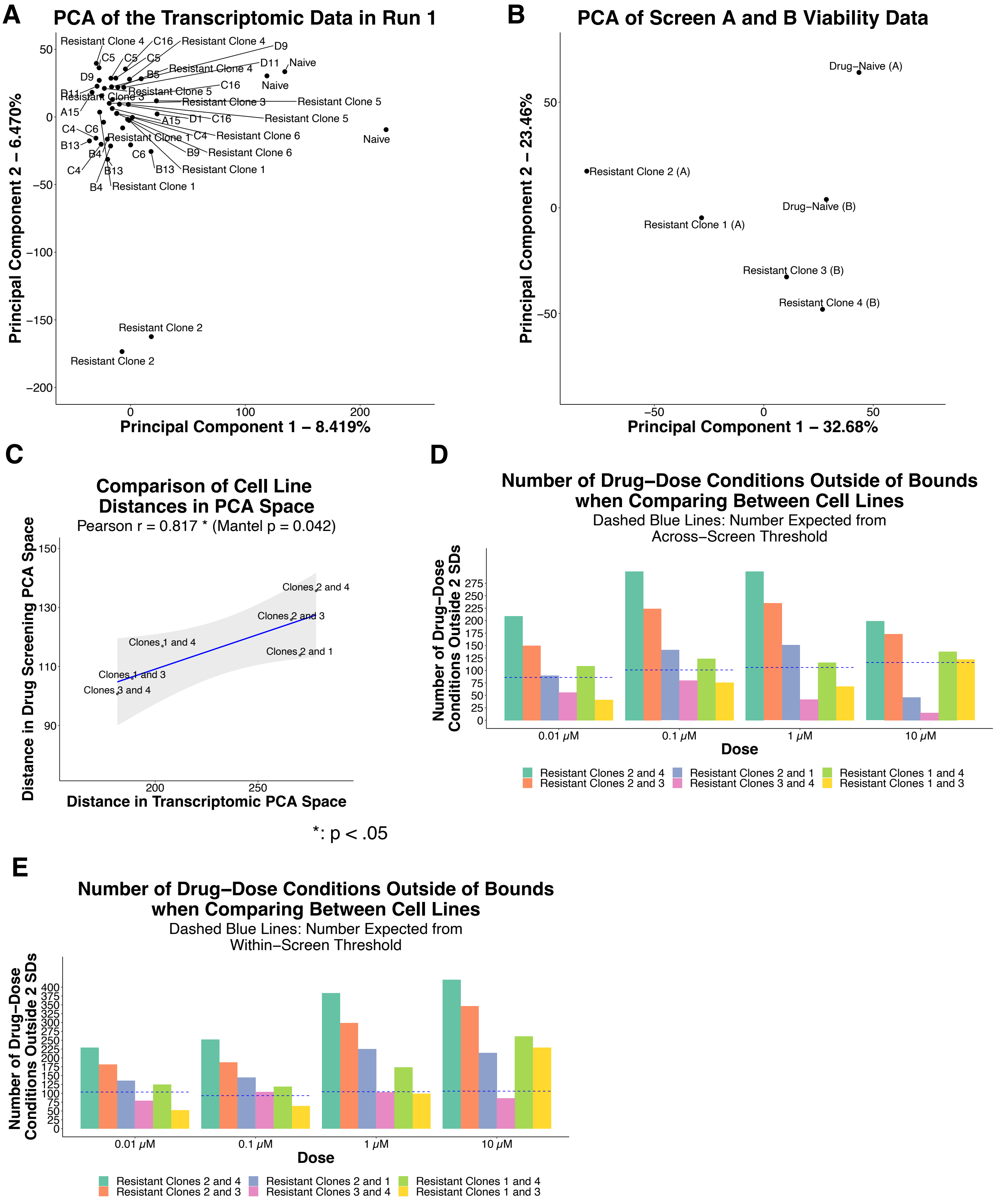

**Supplemental Figure 5.**

**A.** Principal component analysis (PCA) of the first bulk RNA sequencing run, which included the drug-naive cells and the four resistant clones used in high-throughput screening. Resistant Clones 1 - 4 are labelled. Resistant Clones 5 and 6, which were used in follow-up screening, are also labelled. Each point is an individual replicate.
**B.** Principal component analysis (PCA) of the viability data of high-throughput Screens A and B. Each point is an individual replicate and is labelled according to which screen it came from.
**C.** Scatter plot of the distance between pairs of resistant clones in high-throughput screening PCA space (y axis) versus in transcriptomic PCA space (x axis). PCA was conducted on the average of all replicates and the distance between each pair of cell lines was calculated in transcriptomic and in high-throughput screening space. The Pearson correlation coefficient was calculated, and the Mantel test was used to assess statistical significance. Both are shown.
**D.** Grouped bar plot of the number of drug-dose conditions with a difference greater than the across-batch confidence threshold between each pair of resistant clones. Dashed blue lines represent the number expected to be outside by chance based on the across-batch comparison. Bars are colored by clone pair.
**E.** Grouped bar plot of the number of drug-dose conditions with a difference greater than the within-batch confidence threshold between each pair of resistant clones. Dashed blue lines represent the number expected to be outside by chance based on the within-batch comparison. Bars are colored by clone pair.

**Supplemental Figure 6.**

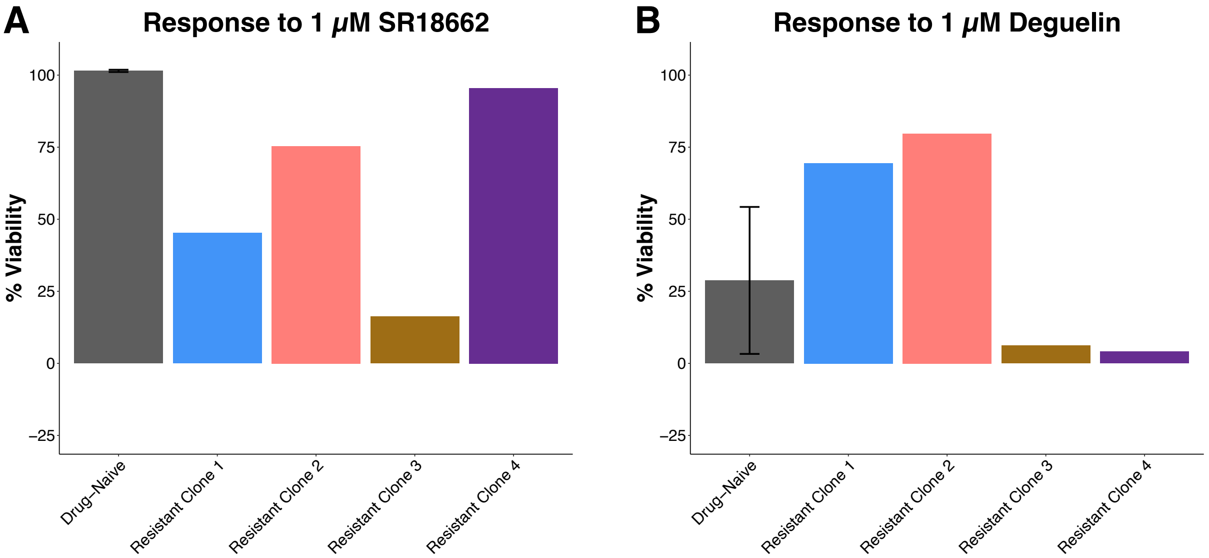


**A.** Bar plot of the % viability of 1 μM SR18662 on each cell line in the high-throughput screens. The bar for the drug-naive cells is the average of the value in each screen. SR18662 is an example of a unique killer because it has less than 30% viability for only one cell line and greater than 30% viability for two cell lines. The values for the resistant clones are from screening with vemurafenib and the value for the drug-naive cells is from screening without vemurafenib.

**B.** Bar plot of the % viability of 1 μM Deguelin on each cell line in the high-throughput screens. The bar for the drug-naive cells is the average of the value in each screen. Deguelin is an example of a shared killer because it has less than 30% viability for two cell lines and greater than 70% viability for two cell lines. The values for the resistant clones are from screening with vemurafenib and the value for the drug-naive cells is from screening without vemurafenib.

### Supplemental Figure 7.

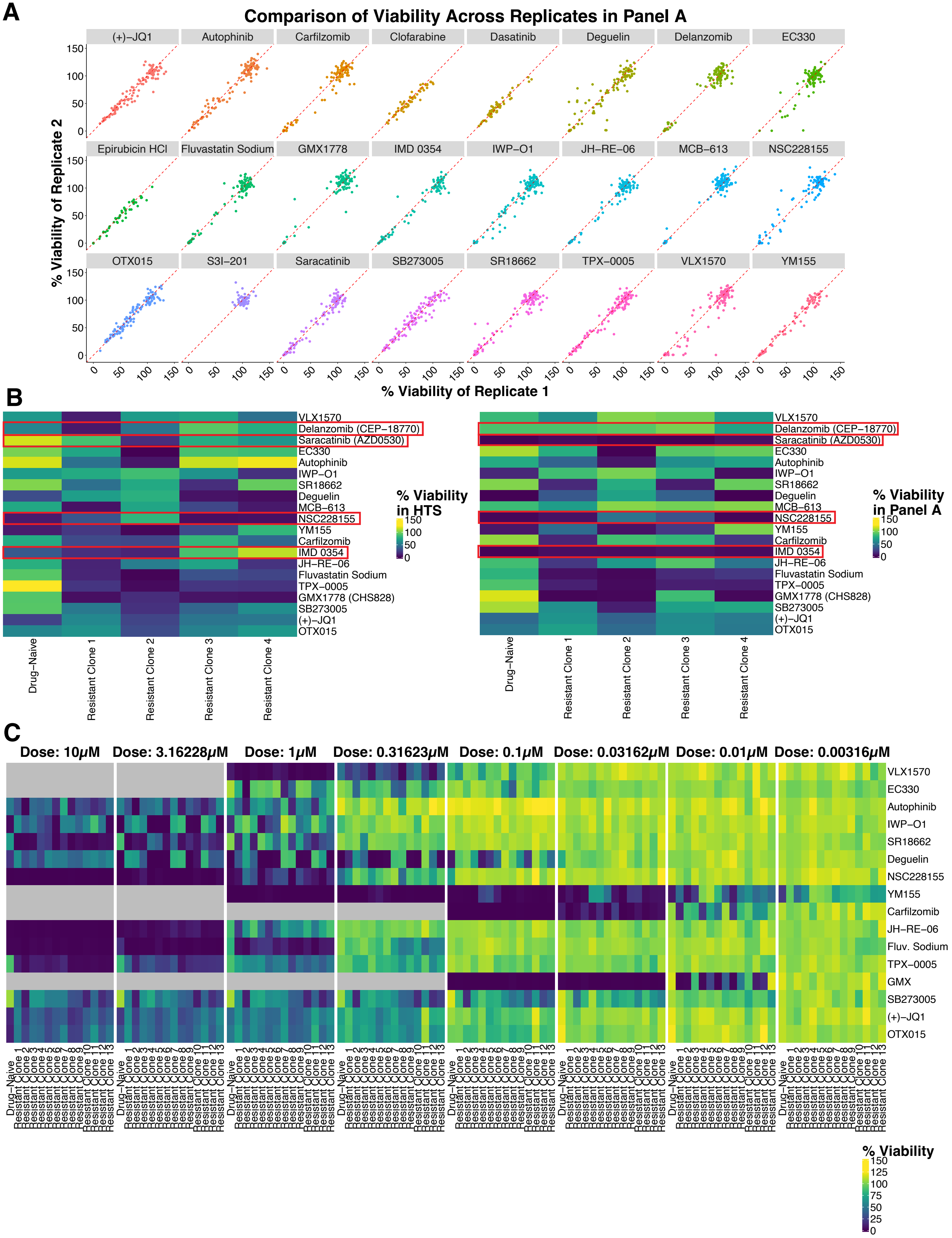

**Supplemental Figure 7.**

**A.** Scatterplots of the viabilities of replicate 2 (y axis) and replicate 1 (x axis) in Follow-Up Panel A. Each point represents a drug-dose condition on a particular cell line. Points are colored by drug and plots are faceted by drug.
**B.** ***(Left)*** Heatmap of the viability of each drug included in Follow-Up Panel A at the dose of interest in the high-throughput screens. The values for the drug-naive cells are the average of the value in each screen. ***(Right)*** Heatmap of the viability of each drug included in Follow-Up Panel A at the dose of interest in Follow-Up Panel A. The values for each cell line are the average of the two replicates in the panel. The viabilities of four drugs (MCB-613, Saracatinib (AZD 0530), IMD 0354, and Delanzomib (CEP-18770)) did not match the viabilities observed in the high-throughput screens at the doses of interest, and thus were not included in further analyses or experiments.
**C.** Heatmaps of the sixteen drugs in Follow-Up Panel A that did validate based on the high-throughput screening data at individual doses. The gray rows indicate drugs that were not tested at that particular dose as the starting dose for that drug was lower. The values for each cell line are the average of the two replicates in the panel.

### Supplemental Figure 8.

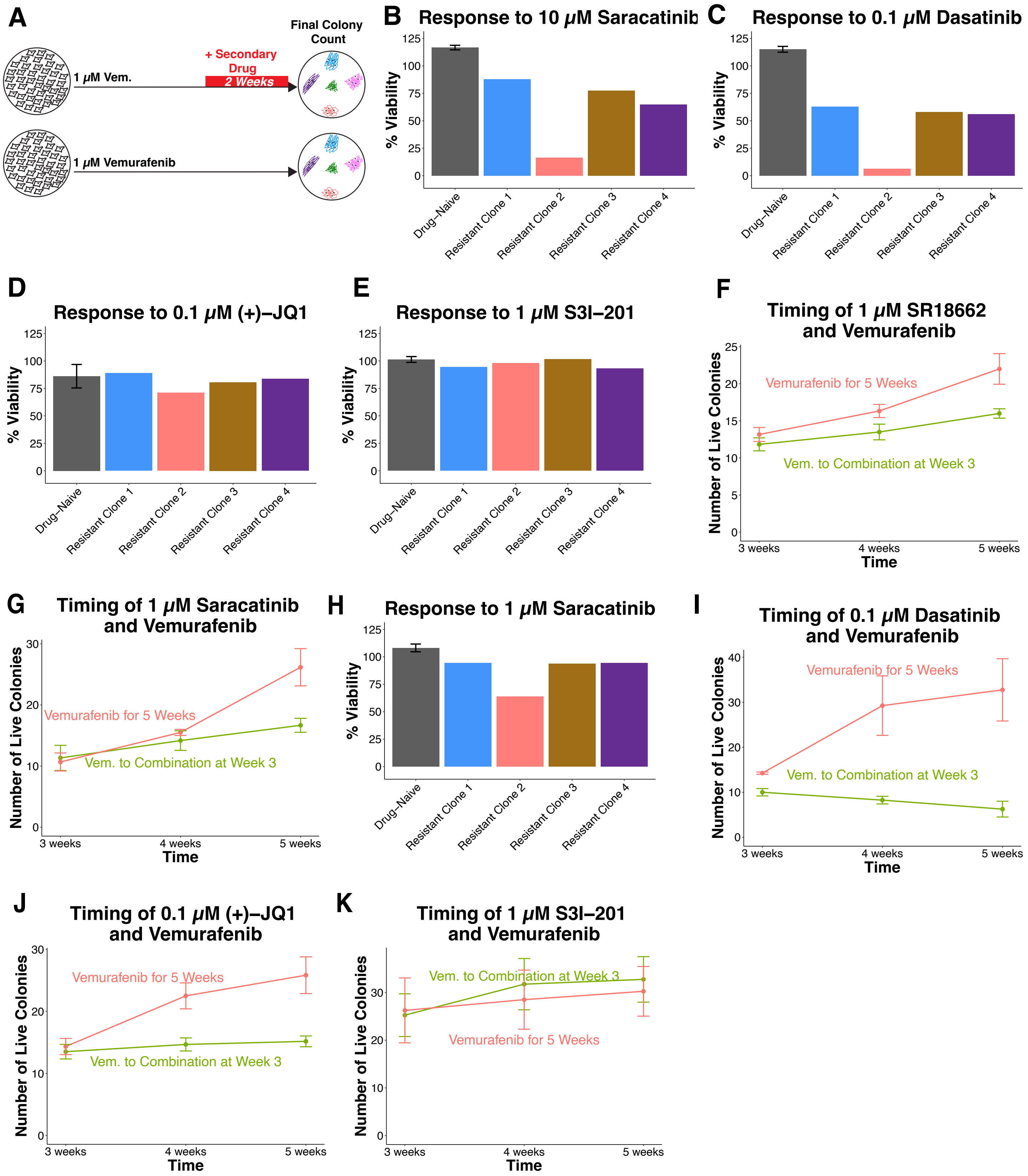

**Supplemental Figure 8.**

**A.** Cartoon schematic showing the experimental design for the experiments. Briefly, drug-naive cells were plated and then treated with either vemurafenib only for five weeks or vemurafenib for three weeks with another drug added for the remaining two weeks. Colony counts were quantified and compared at three, four, and five weeks for all conditions.
**B.** Bar plot of the % viability of 10 $\mu$M saracatinib on each cell line in the high-throughput screens. The bar for the drug-naive cells is the average of the value in each screen. The values for the resistant clones are from screening with vemurafenib and the value for the drug-naive cells is from screening without vemurafenib.
**C.** Bar plot of the % viability of 0.1 $\mu$M dasatinib on each cell line in the high-throughput screens. The bar for the drug-naive cells is the average of the value in each screen. The values for the resistant clones are from screening with vemurafenib and the value for the drug-naive cells is from screening without vemurafenib.
**D.** Bar plot of the % viability of 0.1 $\mu$M (+)-JQ1 on each cell line in the high-throughput screens. The bar for the drug-naive cells is the average of the value in each screen. The values for the resistant clones are from screening with vemurafenib and the value for the drug-naive cells is from screening without vemurafenib.
**E.** Bar plot of the % viability of 1 $\mu$M S3I-201 on each cell line in the high-throughput screens. The bar for the drug-naive cells is the average of the value in each screen. The values for the resistant clones are from screening with vemurafenib and the value for the drug-naive cells is from screening without vemurafenib.
**F.** Line graph of the number of live colonies at three, four, and five weeks after treatment with vemurafenib only or vemurafenib followed by addition of SR18662 after three weeks. The number of live colonies is the average across three experiments, each of which was done in duplicate (n = 6).
**G.** Line graph of the number of live colonies at three, four, and five weeks after treatment with vemurafenib only or vemurafenib followed by addition of 1 $\mu$M saracatinib after three weeks. The number of live colonies is the average across three experiments, each of which was done in duplicate (n = 6, n = 10 for combination treatment).
**H.** Bar plot of the % viability of 1 $\mu$M saracatinib on each cell line in the high-throughput screens. The bar for the drug-naive cells is the average of the value in each screen. The values for the resistant clones are from screening with vemurafenib and the value for the drug-naive cells is from screening without vemurafenib.
**I.** Line graph of the number of live colonies at three, four, and five weeks after treatment with vemurafenib only or vemurafenib followed by addition of dasatinib after three weeks. The number of live colonies is the average across two experiments, each of which was done in duplicate (n = 4).
**J.** Line graph of the number of live colonies at three, four, and five weeks after treatment with vemurafenib only or vemurafenib followed by addition of 0.1 $\mu$M (+)-JQ1 after three weeks. The number of live colonies is the average across three experiments, each of which was done in duplicate (n = 6).
**K.** Line graph of the number of live colonies at three, four, and five weeks after treatment with vemurafenib only or vemurafenib followed by addition of S3I-201 after three weeks. The number of live colonies is the average across two experiments, each of which was done in duplicate (n = 4).

### Supplemental Figure 9.

# A

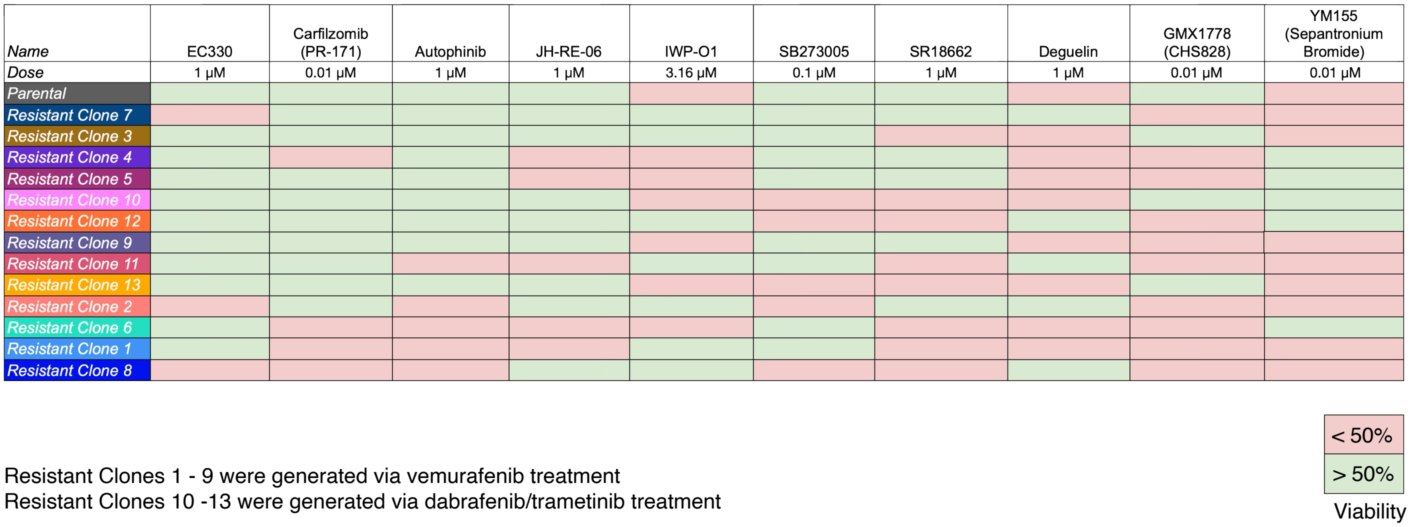


**A.** Chart showing which resistant clones were considered killed (viability less than 50%) or not killed (viability greater than 50%) by each drug at the specified dose in the follow-up panel. Squares colored green represent a clone that was not killed, while red squares represent a clone that was killed.

### Supplemental Figure 10.

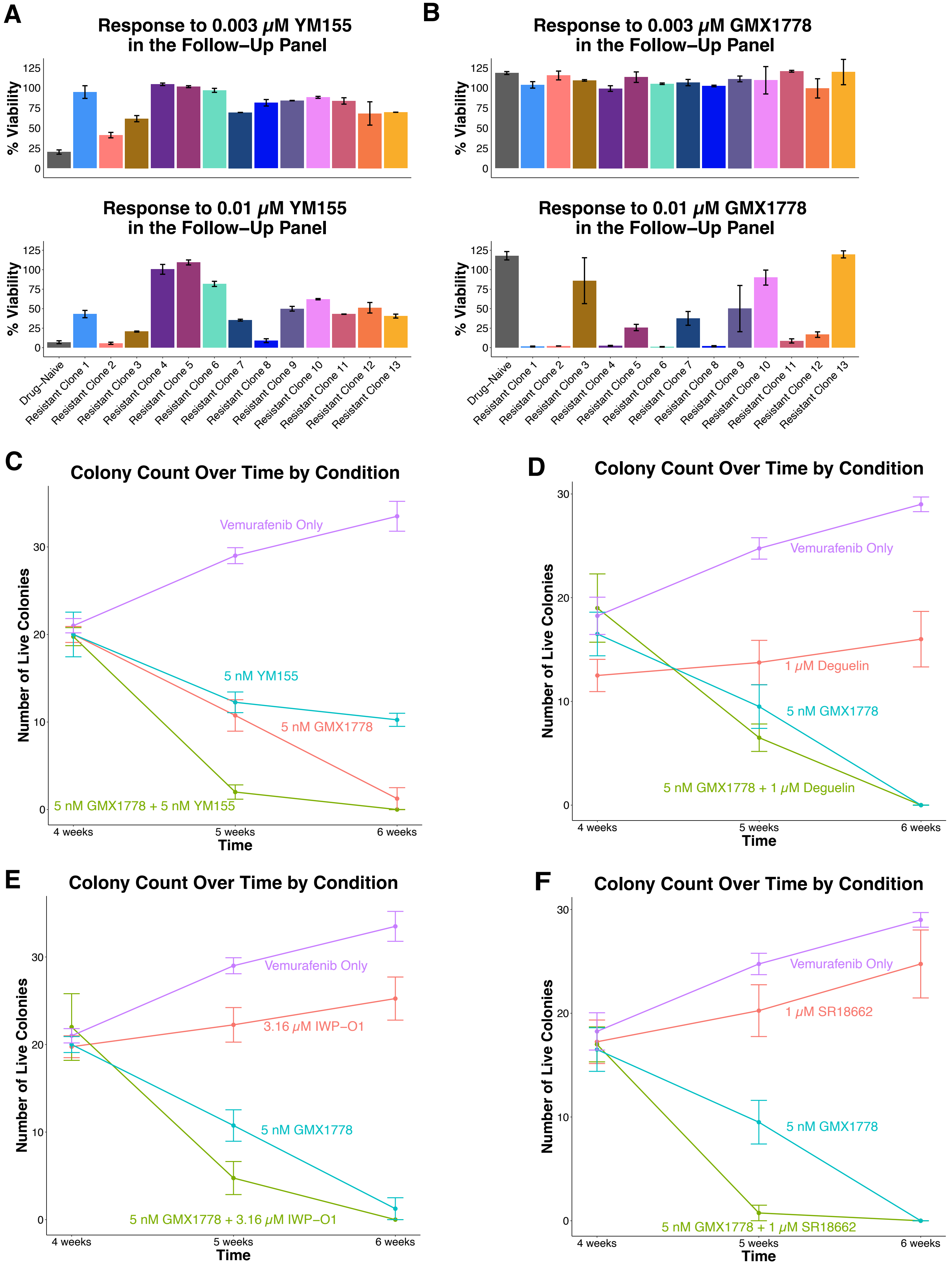

**Supplemental Figure 10.**

**A.** ***(Top)*** Bar graph of the drug-naive cells and resistant clones' responses to 0.003 $\mu$M YM155 in the follow-up panel (seven days of drug treatment). The values for the resistant clones are from screening with vemurafenib (Clones 1 - 9) or dabrafenib/trametinib (Clones 10 - 13). The value for the drug-naive cells is from screening without vemurafenib. All conditions were tested in duplicate, and the average of the value is shown (n = 2). ***(Bottom)*** Bar graph of the drug-naive cells and resistant clones' responses to 0.01 $\mu$M YM155 in the follow-up panel (seven days of drug treatment). The values for the resistant clones are from screening with vemurafenib (Clones 1 - 9) or dabrafenib/trametinib (Clones 10 - 13). The value for the drug-naive cells is from screening without vemurafenib. All conditions were tested in duplicate, and the average of the value is shown (n = 2).
**B.** ***(Top)*** Bar graph of the drug-naive cells and resistant clones' responses to 0.003 $\mu$M GMX1778 in the follow-up panel (seven days of drug treatment). The values for the resistant clones are from screening with vemurafenib (Clones 1 - 9) or dabrafenib/trametinib (Clones 10 - 13). The value for the drug-naive cells is from screening without vemurafenib. All conditions were tested in duplicate, and the average of the value is shown (n = 2). ***(Bottom)*** Bar graph of the drug-naive cells and resistant clones' responses to 0.01 $\mu$M GMX1778 in the follow-up panel (seven days of drug treatment). The values for the resistant clones are from screening with vemurafenib (Clones 1 - 9) or dabrafenib/trametinib (Clones 10 - 13). The value for the drug-naive cells is from screening without vemurafenib. All conditions were tested in duplicate, and the average of the value is shown (n = 2).
**C.** Line graph of the number of live colonies at four, five, and six weeks after treatment with vemurafenib only, vemurafenib with 5 nM GMX1778 added after four weeks, vemurafenib with 5 nM YM155 added after four weeks, or vemurafenib with both drugs added after four weeks. The number of live colonies is the average across two experiments, each of which was done in duplicate (n = 4).
**D.** Line graph of the number of live colonies at four, five, and six weeks after treatment with vemurafenib only, vemurafenib with 5 nM GMX1778 added after four weeks, vemurafenib with 1 $\mu$M deguelin added after four weeks, or vemurafenib with both drugs added after four weeks. The number of live colonies is the average across two experiments, each of which was done in duplicate (n = 4).
**E.** Line graph of the number of live colonies at four, five, and six weeks after treatment with vemurafenib only, vemurafenib with 5 nM GMX1778 added after four weeks, vemurafenib with 3.16 $\mu$M IWP-O1 added after four weeks, or vemurafenib with both drugs added after four weeks. The number of live colonies is the average across two experiments, each of which was done in duplicate (n = 4).
**F.** Line graph of the number of live colonies at four, five, and six weeks after treatment with vemurafenib only, vemurafenib with 5 nM GMX1778 added after four weeks, vemurafenib with 1 $\mu$M SR18662 added after four weeks, or vemurafenib with both drugs added after four weeks. The number of live colonies is the average across two experiments, each of which was done in duplicate (n = 4).

**Supplemental Figure 11.**

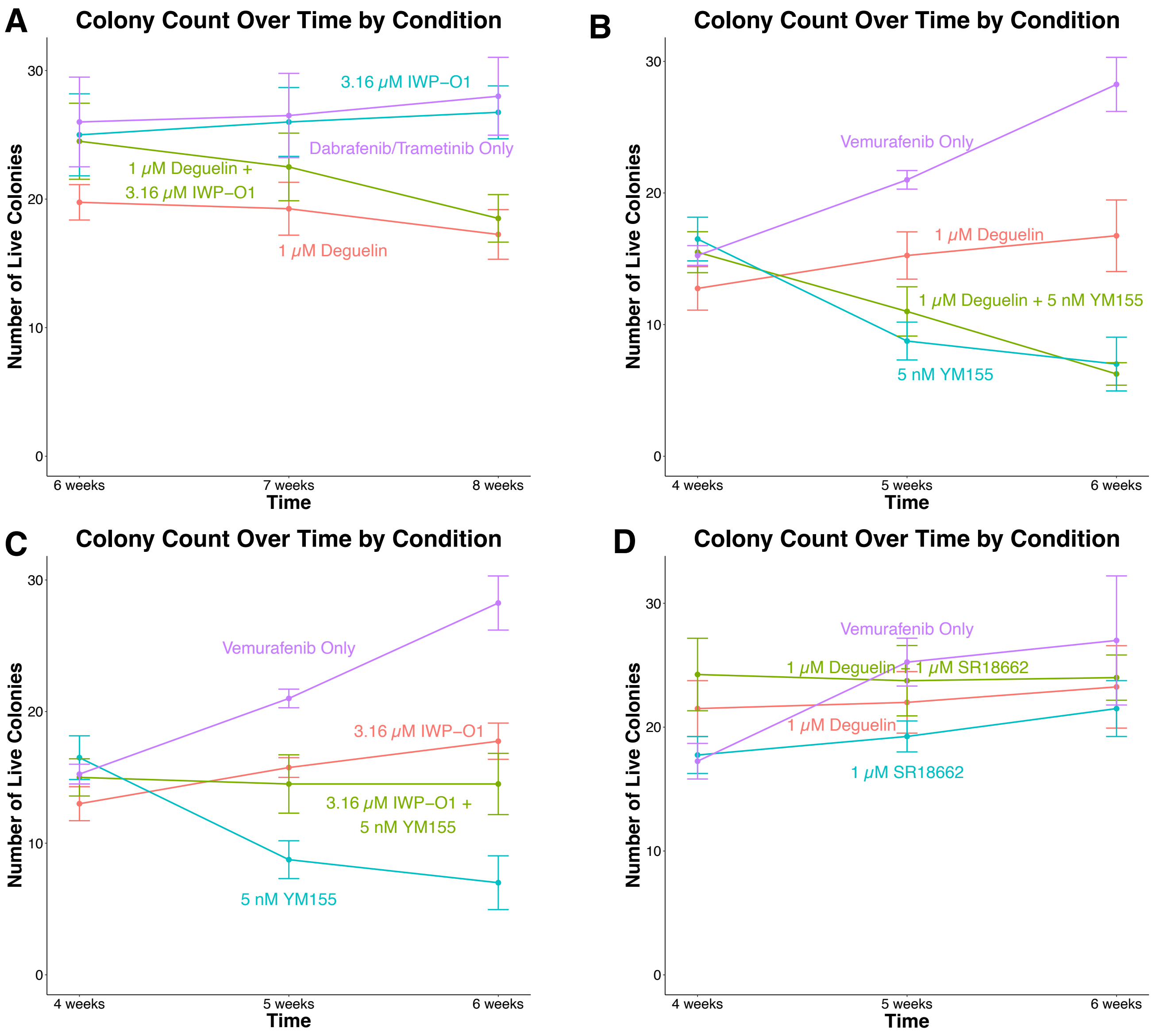


**A.** Line graph of the number of live colonies at four, five, and six weeks after treatment with dabrafenib/trametinib only, dabrafenib/trametinib with either 3.16 μM IWP-O1 or 1 μM deguelin added after four weeks, or dabrafenib/trametinib with both drugs added after four weeks. The number of live colonies is the average across two experiments, each of which was done in duplicate (n = 4).

**B.** Line graph of the number of live colonies at four, five, and six weeks after treatment with vemurafenib only, vemurafenib with 5 nM YM155 or 1 μM deguelin added after four weeks, or vemurafenib with both drugs added after four weeks. The number of live colonies is the average across two experiments, each of which was done in duplicate (n = 4).

**C.** Line graph of the number of live colonies at four, five, and six weeks after treatment with vemurafenib only, vemurafenib with 5 nM YM155 or 3.16 μM IWP-O1 added after four weeks, or vemurafenib with both drugs added after four weeks. The number of live colonies is the average across two experiments, each of which was done in duplicate (n = 4).

**D.** Line graph of the number of live colonies at four, five, and six weeks after treatment with vemurafenib only, vemurafenib with 1 μM SR18662 or 1 μM deguelin added after four weeks, or vemurafenib with both drugs added after four weeks. The number of live colonies is the average across two experiments, each of which was done in duplicate (n = 4).

**Supplemental Figure 12.**

A

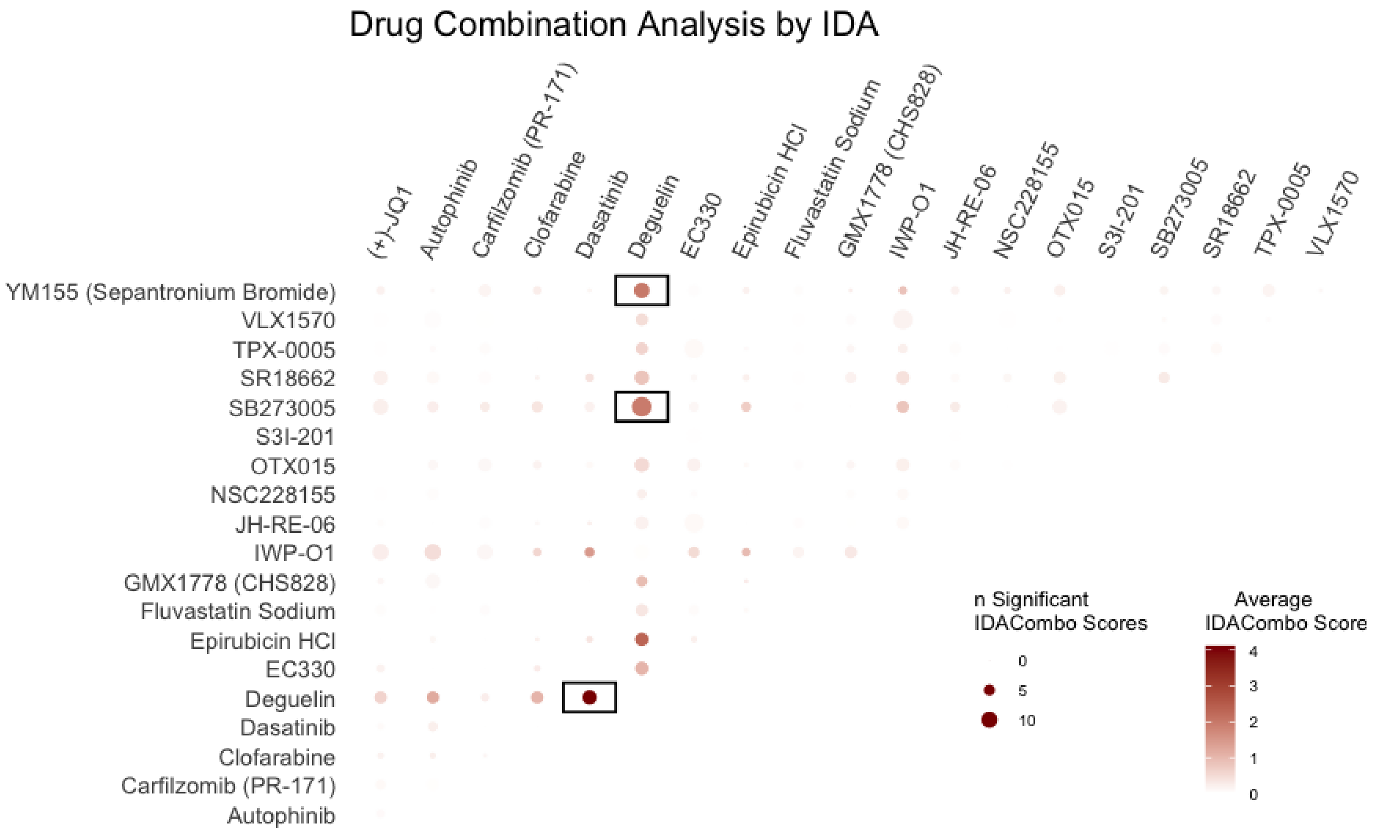


B

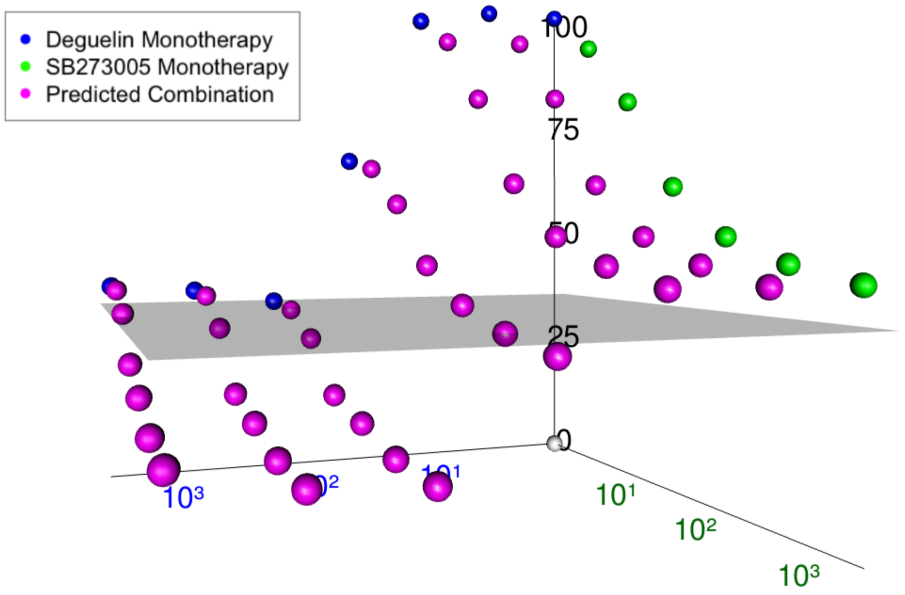


**A.** Bubble chart of the average IDACombo scores of all hypothetical drug combinations based on the follow-up panel viability data, including the drugs that validated based on the high-throughput screening data and the control drugs. Scores were averaged across all doses. Combinations were highlighted if the average IDACombo score was greater than 1 and they were significant across at least 8 dose combinations.
**B.** 3-D plot of experimental cell viability at increasing concentrations of deguelin and SB273005, as well as predicted cell viability at combinations of the two drugs.

### Supplemental Figure 13.

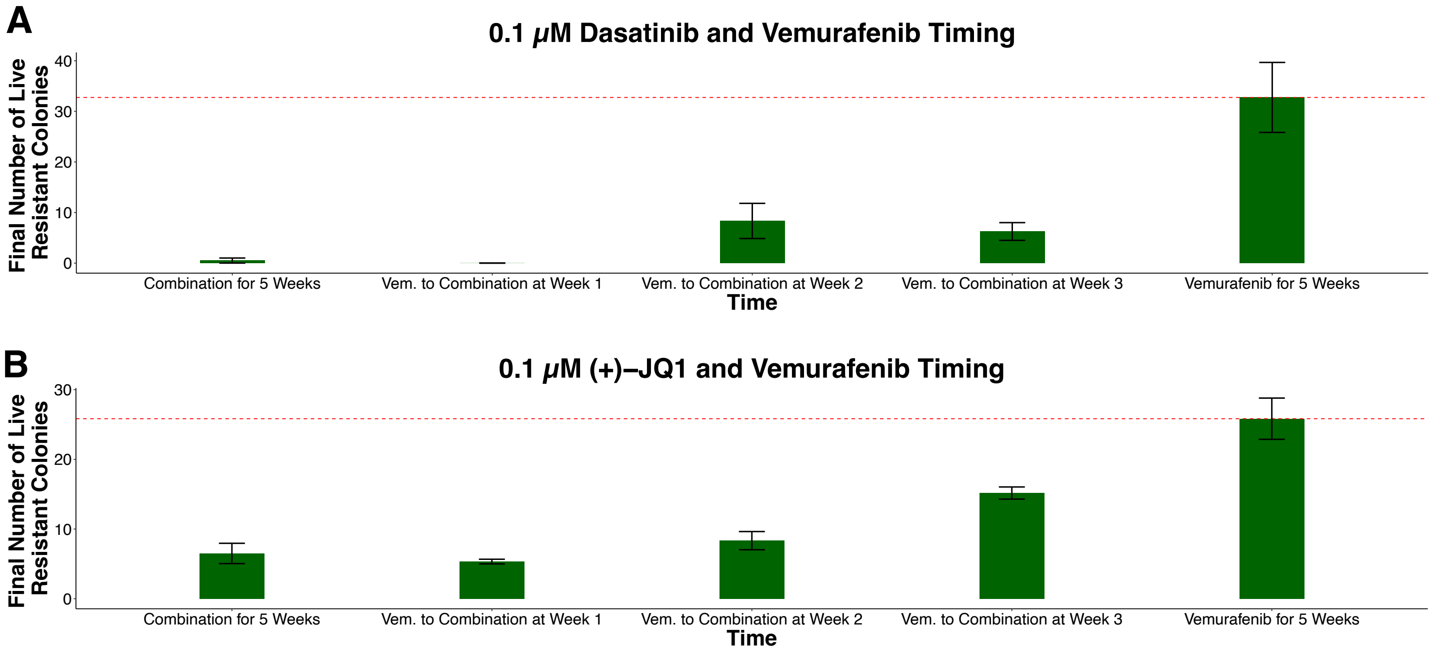


**A.** Bar graph of the number of live colonies at five weeks after treatment with vemurafenib only, vemurafenib and dasatinib, or vemurafenib followed by addition of dasatinib after one, two, or three weeks. The number of live colonies is the average across two experiments, eachof which was done in duplicate (n = 4).

**B.** Bar graph of the number of live colonies at five weeks after treatment with vemurafenib only, vemurafenib and (+)-JQ1, or vemurafenib followed by addition of (+)-JQ1 after one, two, or three weeks. The number of live colonies is the average across three experiments, each of which was done in duplicate (n = 6).
